# Therapeutic scheduling of WEE1 inhibition preserves T cell function and promotes immune control of HPV⁺ tumors

**DOI:** 10.64898/2026.03.10.710574

**Authors:** Ying Liu, Zhiheng Zhang, Yanshan Tao, Lovely Rahgav, Sophie Gray-Gaillard, Yasser Hussaini, Meijie Pan, Justine Shamber, Jeff Kwak, Simone L. Park, Jessica Cramer, Regina Stoltz, Joseph Patria, Jherek Swanger, Kathryn Liu, Malay Kumar Sannigrahi, A. McGarry Houghton, Cristina P. Rodriguez, Ryan M. Carey, Robert Brody, Karthik Rajasekaran, Gregory Weinstein, Gerald P. Linette, Beatriz M. Carreno, Mark M. Painter, E. John Wherry, Bruce Clurman, Devraj Basu, Ahmed Diab

## Abstract

Human papillomavirus–associated oropharyngeal squamous cell carcinoma (HPV⁺ OPC) is driven by viral E6 and E7 oncoproteins, which disrupt G1 checkpoint control and impose selective dependency on WEE1-mediated G2/M regulation. While this vulnerability confers sensitivity to WEE1 inhibition, its immunologic consequences remain poorly defined, and the challenge of eliciting antitumor immunity without compromising immune fitness has limited clinical translation. Here, we show that WEE1 inhibition elicits durable antitumor immunity in immunocompetent models of HPV⁺ OPC. Using murine and human preclinical systems, we demonstrate that the WEE1 inhibitor azenosertib (ZN-c3) mediates tumor control through both cell-autonomous cytotoxicity and immune-dependent mechanisms requiring T cells and conventional dendritic cells. Mechanistically, HPV⁺ tumor cells are deficient in STING signaling and fail to mount canonical type I interferon responses. Instead, tumor cell–intrinsic cGAS drives immune activation through STING-competent host cells within the tumor microenvironment, revealing a non-cell-autonomous relay that circumvents viral immune evasion. Intermittent WEE1 inhibition preserves T cell fitness while maintaining antitumor efficacy, and mice achieving complete responses develop immunologic memory capable of rejecting tumor rechallenge. These findings establish intermittent WEE1 inhibition as an immune-permissive therapeutic strategy that enables antigen-specific T cell responses in HPV-driven malignancies and provides a mechanistic rationale for combination with immunotherapy.

## Introduction

Human papillomavirus–associated oropharyngeal squamous cell carcinoma (HPV⁺ OPC) incidence is rising in the United States (*1, 2*). Although many patients are curable, standard chemo-radiotherapy (CRT) causes substantial toxicity and fails a clinically significant subset with recurrence-prone disease (*3*), underscoring the need for therapies that enable safe de-escalation while improving durable tumor control.

HPV-driven carcinogenesis is mediated primarily by the viral E6 and E7 oncoproteins, which inactivate p53 and RB, respectively, disrupting G1 checkpoint control (*4, 5*). This creates dependence on G2–M checkpoint regulation, rendering HPV⁺ cancers sensitive to pharmacologic WEE1 inhibition (WEE1i) (*6–10*). Consistent with this vulnerability, the first-generation WEE1 inhibitor AZD1775 shows activity in HPV⁺ cancers, which we previously linked to elevated FOXM1 activity (*9*).

Beyond cell-autonomous cytotoxicity, WEE1i–induced DNA damage has been linked to antitumor immune responses in solid tumors, in part through activation of the cGAS–STING pathway (*11–13*). However, because HPV infection antagonizes tumor cell–intrinsic STING signaling (*14, 15*), the relevance of this pathway to WEE1i efficacy in HPV⁺ OPC remains unclear.

Despite promising preclinical activity, clinical translation of WEE1 inhibitors has been limited by dose-limiting hematologic toxicities, particularly with AZD1775 (*16, 17*). Emerging evidence suggests that inhibiting WEE1 can directly affect immune cell proliferation and function (*11, 12, 18*), raising unresolved questions about how to elicit antitumor immunity without compromising immune fitness required for durable tumor control.

The next-generation WEE1 inhibitor azenosertib (ZN-c3) has shown favorable antitumor activity with reduced systemic toxicity in patients with advanced solid tumors (*19–21*), yet its immunologic effects remain undefined. Here, we used immunocompetent preclinical models of HPV⁺ OPC to define the immune contributions to WEE1i efficacy, identify dosing strategies that preserve immune fitness, and characterize the effects of WEE1i on T cell function.

## Results

### Antitumor activity of the WEE1 inhibitor ZN-c3 across immunocompetent models of HPV⁺ OPC

Given the sensitivity of HPV⁺ tumors to WEE1i and emerging evidence that WEE1i can modulate antitumor immunity (*6, 9, 11–13*), we evaluated the antitumor activity of ZN-c3 in immunocompetent models of HPV⁺ OPC. Mice were treated with ZN-c3 (60 mg/kg, 3× weekly; **Fig. 1A**) as intermittent dosing previously showed limited growth delay in xenograft models (*21*).

**Figure 1.**
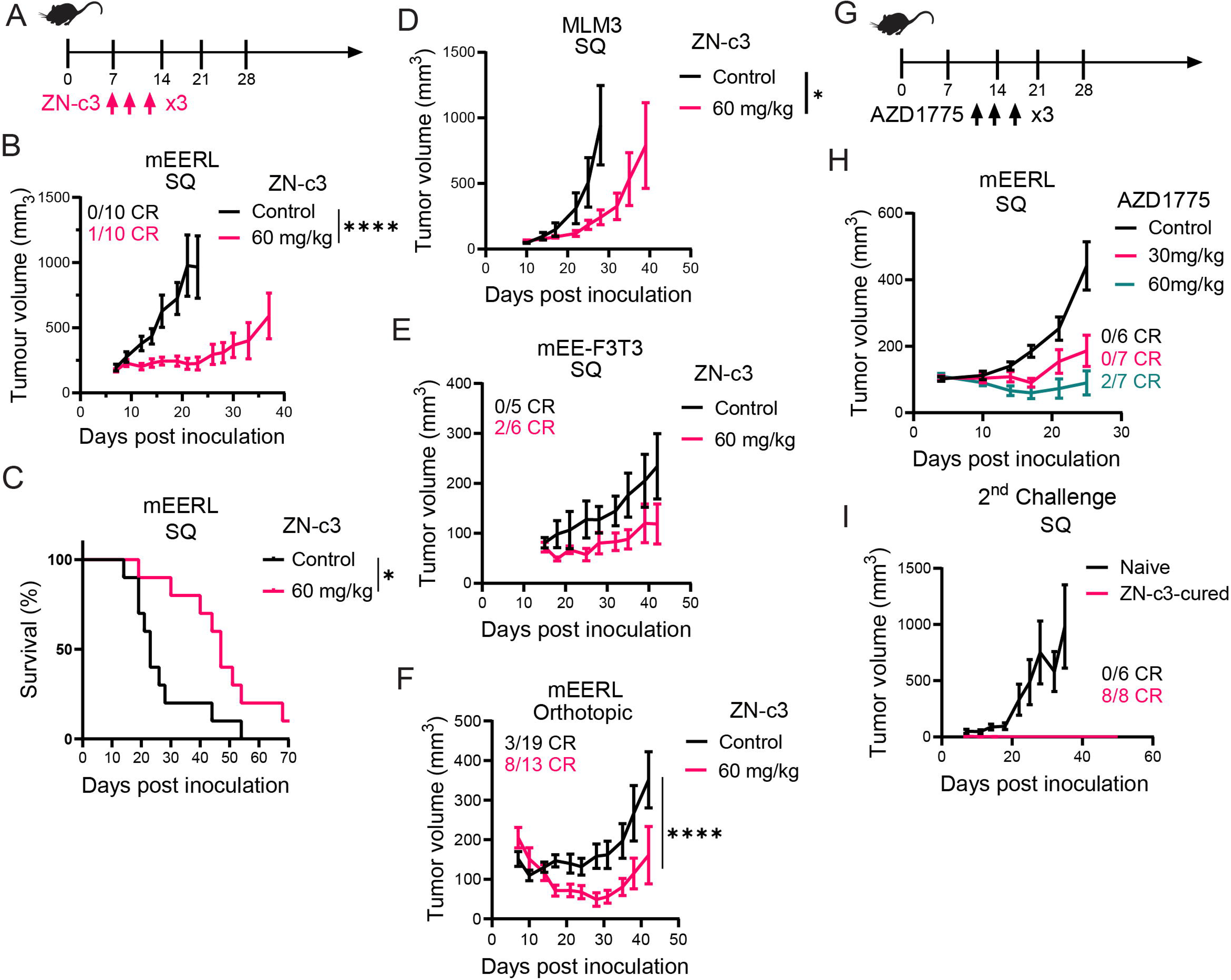
Immunocompetent murine models of HPV^+^ OPC are sensitive to WEE1i. **(A)** Drug administration scheme of the WEE1 inhibitor ZN-c3 (oral gavage). **(B)** Tumor growth curves of WT B6 mice bearing mEERL subcutaneous (SQ) flank tumors, treated with vehicle or 60 mg/kg ZN-c3 as in **A**. Number of mice rejecting the tumor in each group is shown. **(C)** Kaplan–Meier overall survival curves of mice from **B**. **(D)** Tumor growth curves of mice bearing MLM3 flank tumors, treated as in **A**. **(E)** Tumor growth curves of mice bearing mEE-F3T3 flank tumors (HPV16 E6/E7⁺ murine tonsillar epithelial cells expressing FGFR3–TACC3), treated as in **A**. **(F)** Tumor growth curves of mice bearing mEERL orthotopic tumors, treated as in **A**. **(G)** Drug administration scheme of the WEE1 inhibitor AZD1775 (oral gavage). Doses were selected to span submaximal WEE1 inhibition, enabling assessment of immune-dependent versus immune-independent mechanisms of tumor control. **(h)** Tumor growth curves of mice bearing mEERL flank tumors treated as in **F**. **(I)** Tumor growth curves from a rechallenge experiment: Naïve mice and mice cured by prior ZN-c3 treatment were re-inoculated with mEERL cells on the contralateral flank 90 days after rejecting the initial orthotopic tumor. For tumor growth curves, each symbol represents average tumor volume, Error bars represent SEM. Statistical significance was assessed using growth-rate–based tumor modeling, in which individual tumor volumes were fit to an exponential growth model and treatment effects were compared using rate-based metrics with significance indicated (\**, P < 0.05; ****, P < 0.0001*).

We first tested ZN-c3 in the mEERL model, a murine oropharyngeal carcinoma line transformed with HRAS and HPV16 E6/E7 and syngeneic to C57BL/6J (B6) mice (*22*). In the subcutaneous flank setting, ZN-c3 treatment inhibited mEERL tumor growth and prolonged survival of tumor-bearing mice compared with vehicle controls, with 1 of 10 complete responses (CRs) observed (**Fig. 1A–C**). Comparable antitumor activity was observed in MLM3 tumors, a CRT-resistant mEERL variant (*23*), and in mEE-F3T3 flank tumors (*24*), which demonstrated growth suppression including 2 of 6 CRs (**Fig. 1D, E**).

To assess efficacy in an orthotopic context, mEERL cells were implanted into the buccal mucosa of syngeneic hosts. Orthotopic tumors grew more slowly than subcutaneous flank tumors and exhibited spontaneous clearance in a subset of mice (3 of 19; **Fig. 1F**). Intermittent ZN-c3 treatment further suppressed orthotopic tumor growth and increased tumor rejection rates in both mEERL (8 of 13 CRs; **Fig. 1F**) and MLM3 tumors (3 of 6 CRs; **Sup. Fig. 1F**).

To assess target engagement, mice bearing orthotopic mEERL tumors were treated with ZN-c3 or vehicle for six hours. Immunohistochemistry showed reduced pCDK1^Y15^ levels and increased the DNA damage marker γH2AX^S139^ in ZN-c3–treated tumors compared with controls (**Sup. Fig. 1A–C**). Consistent with a class effect of WEE1i, the structurally distinct WEE1 inhibitor AZD1775 resulted in dose-dependent tumor growth suppression in mEERL flank tumors (**Fig. 1G, H**). Intermittent ZN-c3 treatment was well tolerated as assessed by body weight (**Sup. Fig. 1D, E**). Collectively, these results indicate that ZN-c3 monotherapy activity is likely mediated through on-target activity.

ZN-c3 induced durable CRs in a subset of mice (**Fig. 1F; Sup. Fig. 1F**). Mice achieving CR were maintained off treatment and remained tumor-free for more than 90 days prior to mEERL rechallenge. Upon rechallenge, cured mice rejected tumor growth, whereas tumor-naïve controls progressed (**Fig. 1I**), consistent with immunologic memory. Together, these findings indicate that antitumor responses observed across overlapping dosing paradigms and structurally distinct WEE1 inhibitors occur in models with marked WEE1i sensitivity, enabling subsequent dissection of tumor cell–autonomous versus immune-mediated mechanisms.

### Antigen-specific immunity cooperates with WEE1i to promote immune control of HPV^+^ OPC

To distinguish tumor cell–autonomous effects of WEE1i from immune-mediated mechanisms, we compared the activity of low-dose AZD1775 (30 mg/kg, 3× weekly) in mEERL flank tumors grown in immunocompetent B6 mice versus immunodeficient NOD/SCID/IL2Rγ⁻/⁻ (NSG) hosts. This regimen conferred no therapeutic benefit in NSG mice but significantly delayed tumor growth and prolonged survival in B6 mice (**Fig. 2A–C**).

**Figure 2.**
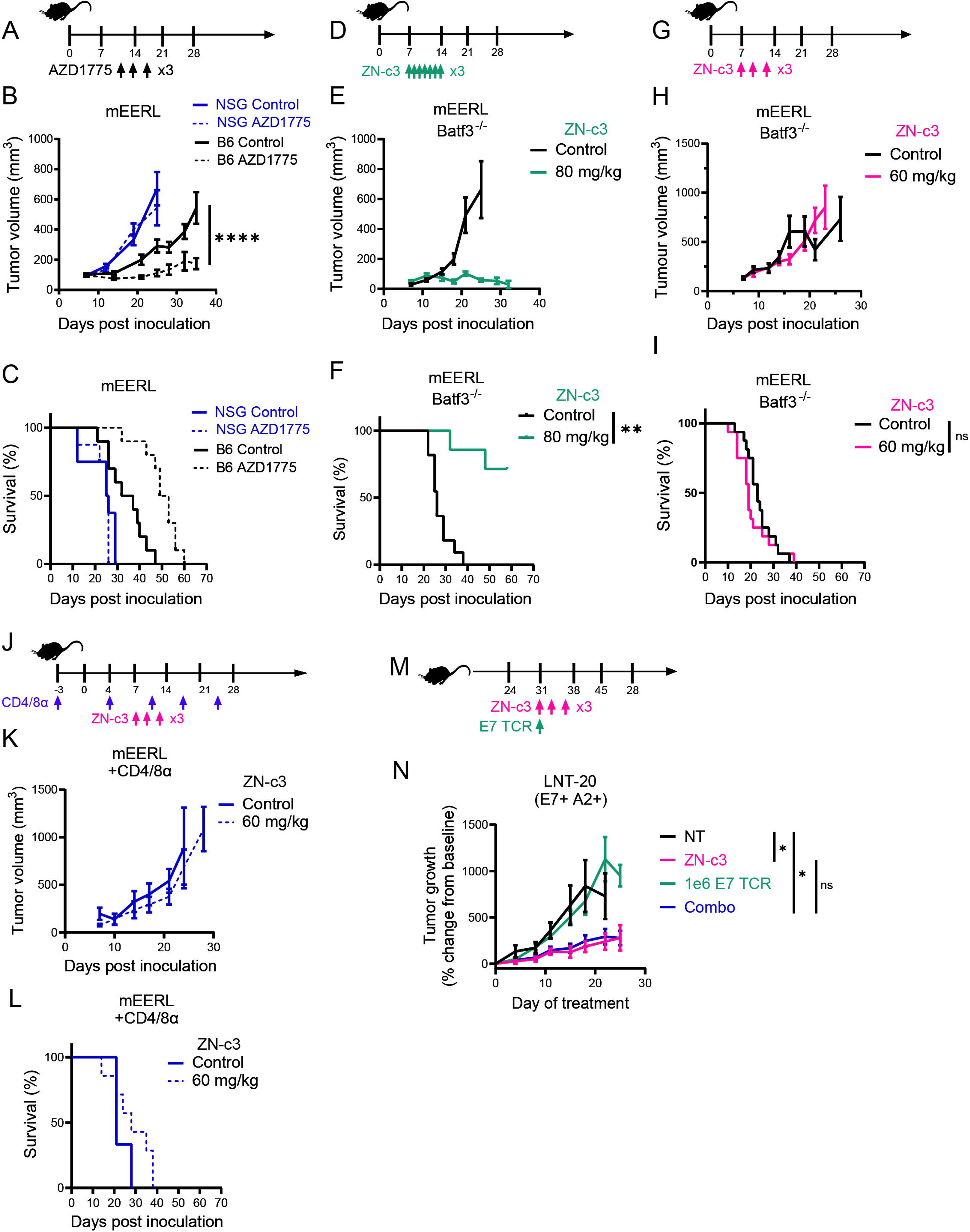
Antigen-specific T cell immunity cooperates with intermittent WEE1i to control HPV⁺ tumors *in vivo*. **(A)** Drug administration scheme for treatment with AZD1775. **(B)** Tumor growth curves of WT B6 and NSG mice bearing established mEERL tumors treated as in **A**. **(C)** Kaplan–Meier overall survival curves for mice from (B). **(D)** Drug administration scheme for treatment with ZN-c3. **(E)** Tumor growth curves of Batf3⁻/⁻ mice bearing established mEERL tumors treated as in **D**. **(F)** Kaplan–Meier survival analysis of mice from (**E**) (P = 0.002). **(G)** Alternative ZN-c3 dosing scheme. **(H)** Tumor growth curves of Batf3⁻/⁻ mice bearing established mEERL tumors treated as in **G**. **(I)** Kaplan–Meier survival analysis of mice from H. **(J)** Experimental schematic for combined CD4⁺ and CD8⁺ T cell depletion and ZN-c3 treatment in WT mice. Administration of anti-CD4 and anti-CD8 antibodies began prior to tumor inoculation (day 0) and continued throughout the experiment. **(K)** Tumor growth curves of WT mice bearing mEERL tumors treated with vehicle or ZN-c3 under conditions of CD4⁺/CD8⁺ T cell depletion, as shown in **J**. **(L)** Kaplan–Meier survival analysis of mice from **K**. **(M)** Treatment scheme for NSG mice bearing HPV16⁺ HLA-A02:01⁺ (HPV⁺ A2⁺) LNT-20 xenografts. Mice received ZN-c3 by oral gavage, HPV E7–specific TCR-engineered T cells (1 × 10⁶ cells, intravenously), or the combination (Combo). Tumor growth was monitored over the indicated treatment period. **(N)** Tumor growth curves of NSG mice from **M**. (\**, P < 0.05; **, P < 0.01; ****, P < 0.0001;* ns, not significant).

To further isolate tumor cell–intrinsic sensitivity to WEE1i, mEERL tumors were established in Batf3^⁻/⁻^ mice, which lack conventional type 1 dendritic cells (cDC1s) required for antigen cross-presentation (*25*). A continuous ZN-c3 regimen (80 mg/kg, 7× weekly) induced durable CRs in 5 of 7 mice (**Fig. 2D–F**), consistent with intrinsic tumor sensitivity to WEE1i. In contrast, the intermittent ZN-c3 regimen (60 mg/kg, 3× weekly), which was highly effective in immunocompetent B6 mice, showed no activity in Batf3^⁻/⁻^ hosts (**Fig. 2G–I**).

To directly assess the contribution of T cells, B6 mice bearing mEERL tumors were treated with intermittent ZN-c3 (60 mg/kg, 3× weekly) in the setting of antibody-mediated T cell depletion (**Sup. Fig. 2A–D**). T cell depletion abrogated the therapeutic benefit conferred by intermittent ZN-c3 (**Fig. 2J–L**). Depletion of either CD8⁺ or CD4⁺ T cells alone partially blunted therapeutic efficacy, indicating nonredundant contributions from both T cell compartments (**Sup. Fig. 2E–H**).

To assess whether antigen-specific human T cells can participate in WEE1i–mediated tumor control, we employed the HPV16⁺ HLA-A*02:01⁺ LNT-20 patient-derived xenograft (PDX) model. NSG mice bearing LNT-20 tumors received adoptive transfer of human CD8⁺ T cells transduced with an HLA-A*02:01–restricted E7–specific TCR. E7 TCR T cells induced dose-dependent tumor regression (**Sup. Fig. 2I**). Intermittent ZN-c3 treatment delayed tumor growth to a similar extent in the presence or absence of E7-specific T cells (**Fig. 2M, N**), indicating that adoptively transferred CD8⁺ T cells do not further augment WEE1i-mediated tumor control in this PDX setting. Together, these findings suggest that additional host-derived immune interactions are likely required to support the WEE1i-induced antitumor immunity.

### Tumor cell-intrinsic cGAS and host STING are required for WEE1i-induced antitumor immunity

WEE1i has been reported to elicit antitumor immune responses in solid tumors through activation of cGAS–STING signaling following DNA damage (*11–13*). Because HPV16 antagonizes tumor cell–intrinsic STING signaling (*14, 15*), we sought to define the mechanisms by which ZN-c3 elicits adaptive immunity in mEERL tumors.

mEERL cells or isogenic non–HPV-transformed mouse oropharyngeal epithelial (mTE) cells were treated with ZN-c3 for 24 hours followed by drug washout. WEE1i reduced CDK1^Y15^ phosphorylation in both cell lines in a dose-dependent and reversible manner (**Fig. 3A–C**). HPV16 E6/E7–transformed mEERL cells accumulated substantially higher levels of DNA damage and premature mitosis compared with mTE controls, and DNA damage in mEERL cells persisted following washout (**Fig. 3A, B, D; Sup. Fig. 3A**), consistent with our previous observations in human HPV⁺ models of OPC (*9*). In contrast, minimal residual DNA damage was detected in mTE cells after recovery (**Fig. 3D**).

**Figure 3.**
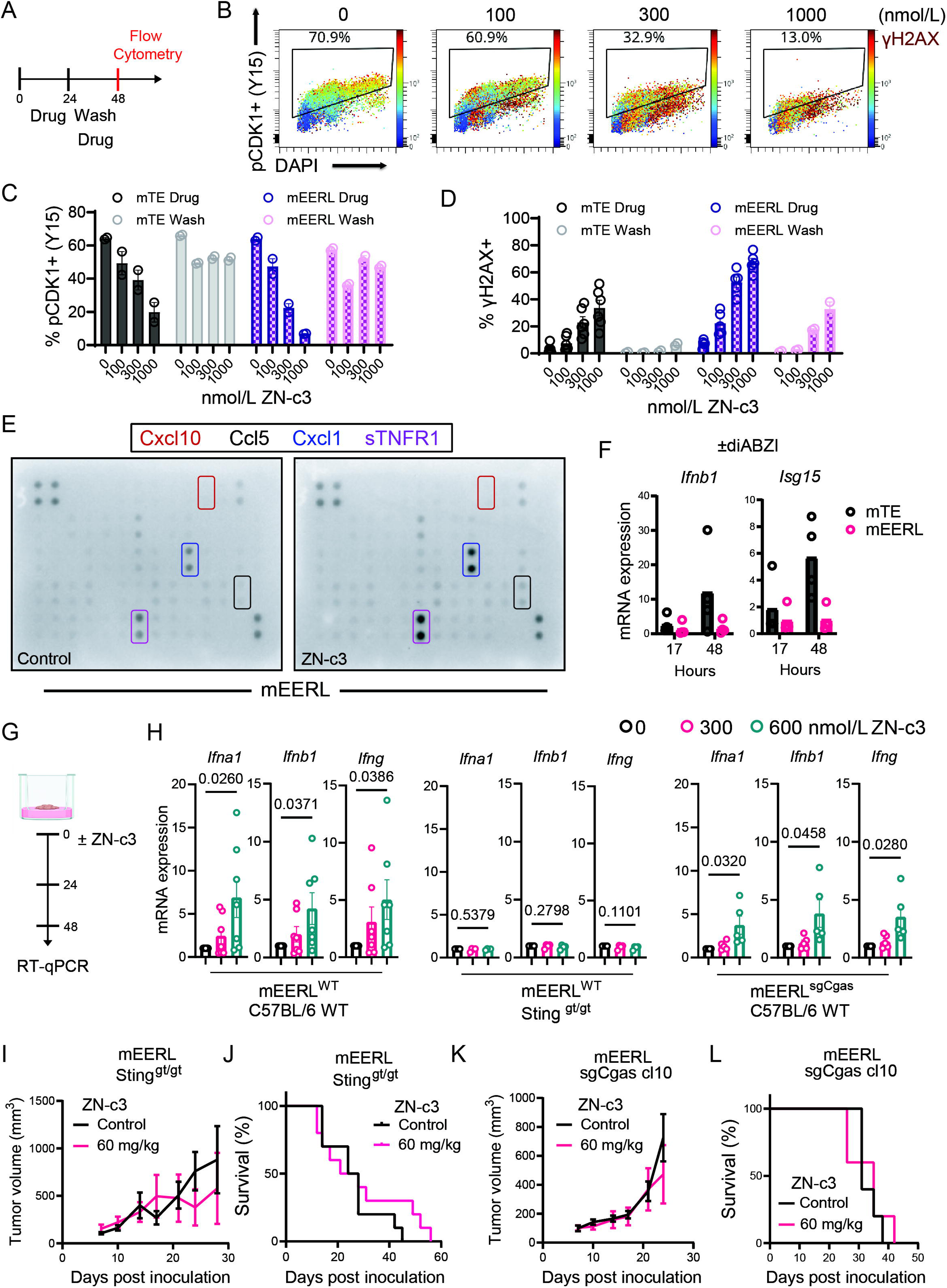
Tumor cell cGAS and host STING contribute to the antitumor immunity triggered by WEE1i. **(A)** Drug treatment scheme for mEERL cells. **(B)** Representative flow cytometry plots showing pCDK1^Y15^ and γH2AX^S139^ levels in mEERL cells treated with ZN-c3 for 24 hours. Flow cytometric quantification of pCDK1^Y15^ **(C)** and γH2AX^S139^ levels **(D)** in isogenic mTE and mEERL cells following ZN-c3 treatment and subsequent drug washout as in **A**. **(E)** Chemokine and cytokine profiling of mEERL cell lysates and culture supernatants following treatment with ZN-c3 (500 nmol/L) or vehicle for 72 hours. Highlighted analytes include Cxcl10 (red), Cxcl1 (blue), Ccl5 (black), and soluble Tnfr1 (purple). **(F)** mTE and mEERL cells were treated with 1 μmol/L diABZI, a STING agonist. Relative mRNA expression of *Ifnb* and *Isg15* was measured by RT-qPCR at 17 and 48 hours post-treatment. **(G)** Drug treatment scheme for *ex vivo* tumor slice cultures generated from mEERL tumors **(H)** Tumor slice cultures generated from wild-type (WT) mEERL tumors grown in B6 WT or *Sting*^gt/gt^ mice, as well as WT B6 mice bearing cGAS-deficient (sg_cGas)mEERL tumors, were treated with ZN-c3 as in **(G)**. Relative mRNA expression of *Ifna1*, *Ifnb1*, and *Ifng* was assessed by RT-qPCR. Statistical significance was determined using unpaired t-test comparing 600 nmol/L ZN-c3 to vehicle group. **(I)** *Sting*^gt/gt^ mice bearing mEERL flank tumors were treated with vehicle or ZN-c3 (60 mg/kg). Tumor growth curves are shown. **(J)** Kaplan–Meier survival analysis of mice from **(H)**. **(K)** WT mice bearing cGAS-deficient mEERL tumors (sg_cGas, clone #10) were treated with vehicle or ZN-c3 (60 mg/kg). Tumor growth curves are shown. **(L)** Kaplan–Meier survival analysis of mice from **(K)**.

Despite robust DNA damage, inhibiting WEE1 using ZN-c3 or AZD1775 failed to induce expression of Cxcl10, Ccl5, type I interferon (IFN) transcripts, STAT1 activation, or interferon-stimulated genes (ISGs) in mEERL cells *in vitro* (**Fig. 3E; Sup. Fig. 3B, C**). Instead, ZN-c3 increased expression of CXCL1 and soluble TNFR1 (**Fig. 3E**), factors associated with immunosuppressive signaling (*26*). Poly(I:C), but not a double-stranded DNA analog, activated type I IFN signaling in mEERL cells, whereas the STING agonist diABZI induced *Ifnb1* and *Isg15* expression in mTE but not mEERL cells (**Fig. 3F; Sup. Fig. 3A, B**). Consistent with these findings, ZN-c3 did not induce *CXCL10* or *CD274* expression in human HPV⁺ OPC models *in vitro* (**Sup. Fig. 3D, E**), indicating compromised tumor cell–intrinsic STING signaling across murine and human systems.

In contrast to these tumor cell–intrinsic *in vitro* findings, WEE1i induced a robust IFN response *in vivo*. AZD1775 treatment of mEERL tumors resulted in upregulation of *Ifna1*, *Ifnb1*, *Stat1*, and multiple ISGs, including *Cxcl9*, *Cxcl10*, and *Cd274* (**Sup. Fig. 3F**). NanoString profiling further revealed enrichment of antigen presentation, IFN signaling, and T cell activation pathways within the tumor microenvironment (TME, **Sup. Fig. 3G**).

To determine whether host STING signaling mediates this IFN response, we generated organotypic tumor slice cultures from mEERL tumors grown in WT or STING-deficient (Sting^gt/gt^) hosts. ZN-c3 induced dose-dependent IFN signaling in slices from WT but not Sting^gt/gt^ hosts (Fig. 3G, H). Accordingly, ZN-c3 treatment (60 mg/kg, 3× weekly) inhibited tumor growth and prolonged survival in WT but not STING-deficient mice (**Fig. 3I, J; compare with Fig. 1A-C**).

Finally, to assess whether tumor cell–intrinsic cGAS contributes to WEE1i-induced immunity, we generated cGas-knockout mEERL cells and established slices from cGas-knockout tumors grown in WT hosts (Sup. Fig. 3H). Tumor cell–specific deletion of cGas attenuated the IFN response to ZN-c3 *ex vivo* (**Fig. 3H**) and completely abrogated therapeutic efficacy *in vivo* (**Fig. 3K, L; Sup. Fig. 3I, J**). Together, these data demonstrate that WEE1i-induced antitumor immunity requires tumor cell–intrinsic cGAS to initiate danger signaling and host STING to propagate productive immune activation within the TME.

### WEE1i potentiates antitumor immunity against HPV⁺ OPC tumors by altering the TME

To define how WEE1i alters the TME, we performed FLEX single-cell whole-transcriptome sequencing (scRNA-seq) on dissociated mEERL tumors grown in B6 mice and treated with AZD1775 or vehicle control (**Sup. Fig. 4A**). Using established lineage-specific markers, cells were clustered into three major compartments—immune, stromal, and tumor (**Fig. 4A**). Compartment-specific gene set variation analysis revealed transcriptional programs consistent with enhanced antitumor immunity, including increased IL-2/STAT5 signaling within immune cells and concomitant upregulation of IFN signaling in tumor and stromal compartments (**Fig. 4B**). In parallel, we observed downregulation of proliferation-associated pathways, including E2F targets and KRAS signaling, within tumor and stromal cells (**Fig. 4B**). We did not detect evidence of integrated stress response (ISR) activation under the AZD1775 dosing regimen used in this experiment (**Sup. Fig. 4C**).

**Figure 4.**
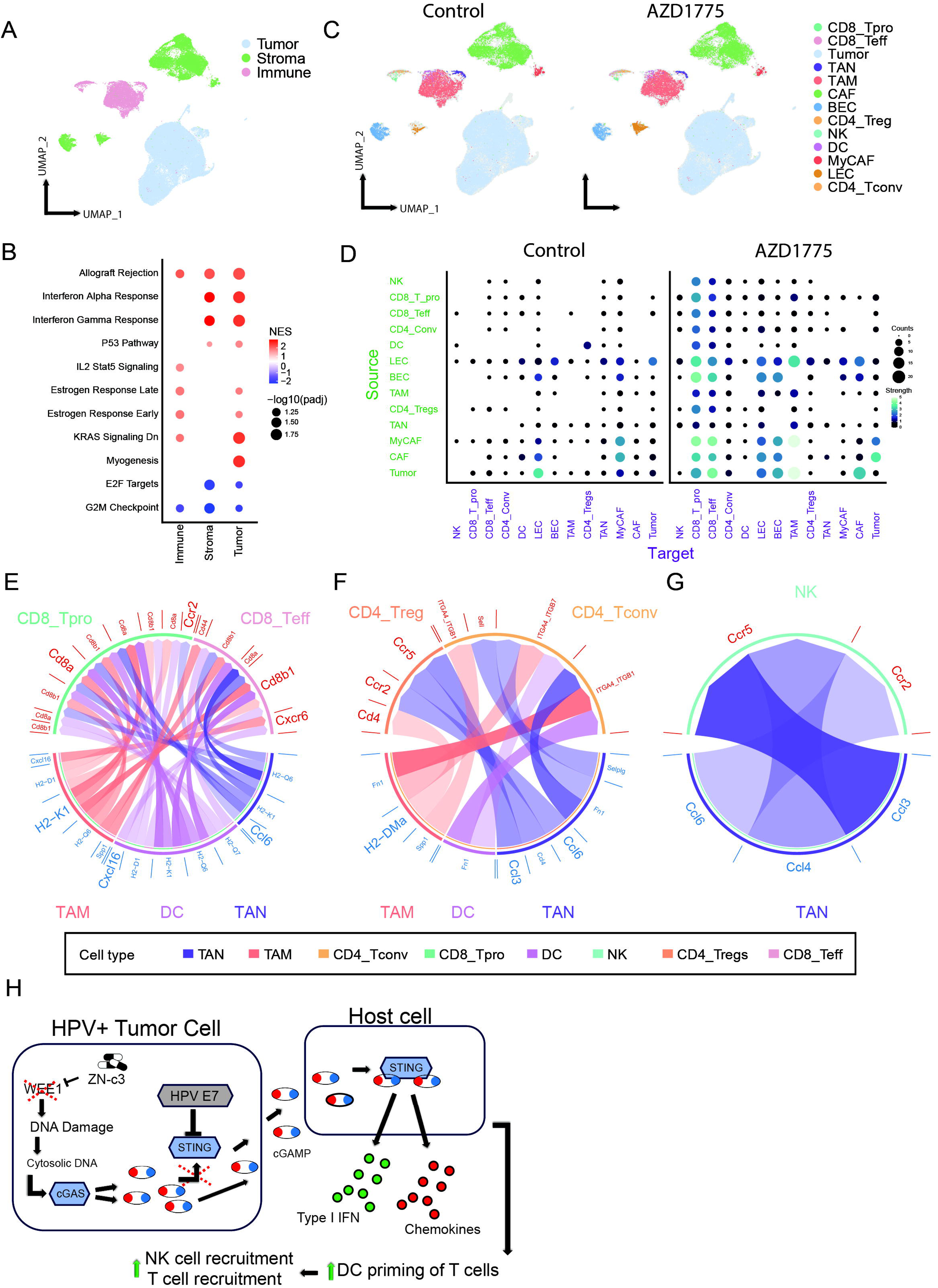
WEE1i reshapes the TME of HPV⁺ mEERL tumors to support lymphocyte recruitment and maintenance. mEERL tumors from vehicle- or AZD1775-treated B6 mice (60 mg/kg, 4 doses over 8 days, n = 4 per group) were dissociated into single cells, barcoded and pooled for single-cell RNA sequencing (scRNA-seq), and subsequently demultiplexed by treatment condition. **(A)** Uniform Manifold Approximation and Projection (UMAP) of scRNA-seq profiles colored by major cellular compartments: tumor, stromal, and immune. **(B)** Compartment-specific Hallmark gene set enrichment analysis (GSEA) based on differentially expressed genes between AZD1775-treated and vehicle-treated tumors. Normalized enrichment scores (NES) are shown for significantly altered pathways, with significance determined using the Benjamini–Hochberg false discovery rate correction. **(C)** Treatment-specific UMAPs colored by annotated cellular subclusters. **(D)** CellChat-based inference of cell–cell communication networks across annotated subclusters. Bubble plots summarize treatment-specific interaction numbers and interaction strength between source and target populations. **(E–G)** Circos plots depicting the top ligand–receptor interactions predicted by CellChat to be enriched following AZD1775 treatment. Connection transparency reflects inferred interaction strength. Antigen-presenting cell (APC) populations—including TAMs, TANs, and DCs—are shown as source populations, with lymphocyte subsets shown as targets: **(E)** CD8⁺ T cells, **(F)** CD4⁺ T cells, and **(G)** NK cells. Abbreviations: CD8_Tpro, CD8⁺ progenitor-like T cells; CD8_Teff, CD8⁺ effector T cells; CD4_Tconv, conventional CD4⁺ T cells; CD4_Tregs, regulatory CD4⁺ T cells; NK, natural killer cells; DC, dendritic cells; TAN, tumor-associated neutrophils; TAM, tumor-associated macrophages; CAF, cancer-associated fibroblasts; MyCAF, myofibroblastic CAFs; BEC, blood endothelial cells; LEC, lymphatic endothelial cells. **(H)** DNA damage induced by WEE1 inhibition engages tumor cell cGAS and TME STING to promote a type I IFN response leading to cDC1-mediated T cell infiltration and immune clearance.

To further resolve TME dynamics, we subclustered cells into 13 populations encompassing endothelial cells, fibroblasts, tumor-infiltrating lymphocytes (TILs), myeloid cells, and tumor cells (**Fig. 4C; Sup. Fig. 4B**). Within the immune compartment, we observed a significant increase in the proportion of effector CD8⁺ TILs following AZD1775 treatment (**Sup. Fig. 4D, F**). These findings were corroborated by multiplex immunohistochemistry (mIHC) staining for pan-cytokeratin, CD8, and Ly6G, which demonstrated an increased CD8:neutrophil ratio in AZD1775-treated tumors compared with vehicle controls (**Sup. Fig. 4G, H**). Analysis of the stromal compartment revealed an approximately fivefold expansion of lymphatic endothelial cells (LECs), accompanied by a reduction in immunosuppressive myofibroblastic cancer-associated fibroblasts (MyCAFs; **Fig. 4C; Sup. Fig. 4E**), consistent with a TME permissive to antitumor immune activity.

To better understand AZD1775–associated signaling within the TME, we performed cell–cell interaction analysis using the CellChat framework (*27*). Intermittent AZD1775 treatment profoundly reshaped ligand–receptor communication networks, increasing both the frequency and strength of inferred interactions (715 AZD1775-specific interactions versus 255 interactions in vehicle-treated tumors), with the most pronounced effects observed within the immune compartment (**Fig. 4D**). Focusing on antigen-presenting cell (APC) populations—including DCs, tumor-associated macrophages (TAMs), and tumor-associated neutrophils (TANs)—as sender populations and CD4⁺ TILs, CD8⁺ TILs, and NK cells as receiver populations, CellChat identified several ligand–receptor programs selectively enriched in AZD1775-treated tumors. Prominent among these was enhanced CD8 receptor signaling through MHC class I expressed by APCs (**Fig. 4E**), consistent with increased antigen presentation and TIL activation. CellChat also predicted enhanced CXCR6–CXCL16 signaling between CD8⁺ TILs and TAMs/DCs (**Fig. 4E**), a pathway implicated in tissue localization of T cells, including tissue-resident memory and tissue-resident exhausted T cells, and associated with enhanced cytotoxic function *in vivo* (*28, 29*). Additionally, increased activity of the CCL3/4/6–CCR2/5 axis between neutrophils and lymphocytes in AZD1775-treated tumors is consistent with enhanced lymphocyte recruitment and persistence (**Fig. 4E-G**).

To define how WEE1i modulates antitumor immune states within the local TME in a human-relevant setting, we first analyzed immune responses in organotypic tumor slice cultures derived from treatment-naïve HPV⁺ OPC specimens. To confirm if TILs can be activated in this *ex vivo* system, tumor slices were stimulated with PMA (phorbol 12-myristate 13-acetate) and ionomycin, followed by dissociation and flow cytometric immunophenotyping. This stimulation induced robust CD8⁺ TIL activation, marked by increased expression of CD25 and CD38, concomitant downregulation of memory-associated markers CD127 and TCF1, and upregulation of inhibitory receptors including CTLA-4 and LAG-3 (**Sup. Fig. 6A**).

Tumor slices were subsequently treated with ZN-c3 or vehicle control, and immunohistochemical analysis confirmed on-target drug activity (**Sup. Fig. 6B, C**). Following dissociation, unsupervised hierarchical clustering of TILs revealed heterogeneous responses to ZN-c3, including an effector-like subset characterized by increased CD25, granzyme B, and CD38 expression, a second subset enriched for memory-associated features (CD127, TCF1), and a third subset with minimal phenotypic change (**Fig. 5A–D**). To explore factors underlying this heterogeneity, we examined baseline immune features associated with divergent responses and found that increased baseline frequencies of activated (CD38⁺) CD4⁺ TILs, including both regulatory and conventional subsets, were negatively associated with induction of a stem-like T cell phenotype following ZN-c3 treatment (**Sup. Fig. 6D, E**).

**Figure 5.**
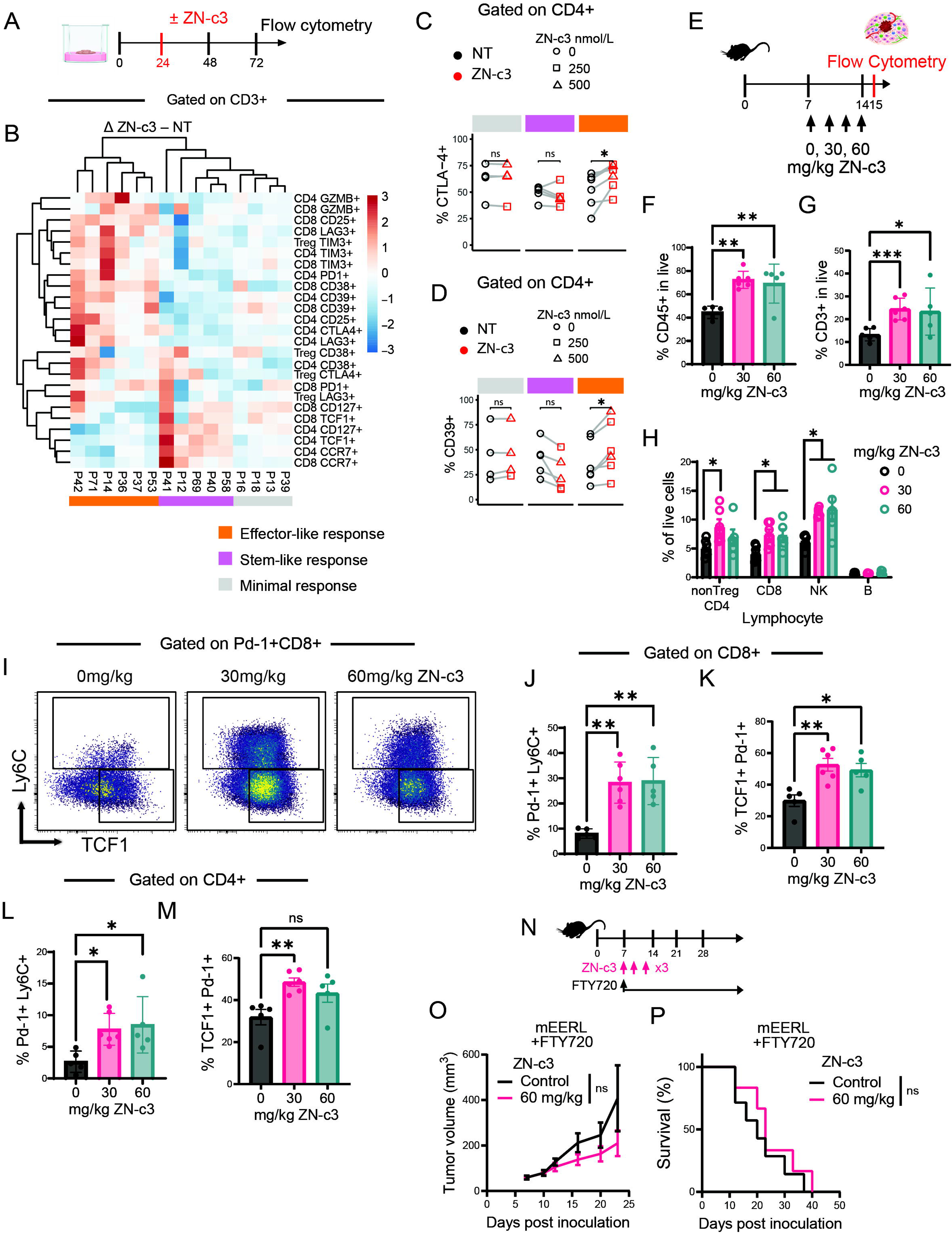
ZN-c3 increases immune infiltration and maintenance of cytotoxic lymphocytes with features of functional persistence. **(A)** Schematic of *ex vivo* ZN-c3 treatment of human tumor slices followed by flow cytometric profiling of TILs. **(B)** Hierarchical clustering heatmap of Z-score–scaled differences in population frequencies for the indicated phenotypic subsets among CD3⁺ TIL following in vitro ZN-c3 treatment relative to untreated controls (Δ ZN-c3 – NT). Rows represent individual immune cell populations and columns represent samples. Color intensity reflects relative enrichment or depletion based on Z-score normalization across populations. Sample groupings corresponding to effector-like, stem-like, or minimal responses are indicated. **(C-D)** Frequency of CTLA-4⁺ (**C**) and CD39⁺ (**D**) cells among non-regulatory CD4⁺ TILs from ex vivo tumor slice culture treated ± ZN-c3 at the indicated concentrations. **(E)** Experimental schematic of the *in vivo* ZN-c3 dosing regimen and timeline for flow cytometric analysis of tumor-infiltrating immune cells. Tumor-bearing mice received ZN-c3 at the indicated doses prior to tumor harvest. **(F)** Frequency of CD45⁺ cells among live tumor cells following treatment with the indicated doses of ZN-c3. **(G)** Frequency of CD3⁺ TILs among live tumor cells following ZN-c3 treatment. **(H)** Proportions of indicated lymphocyte populations among live tumor-infiltrating immune cells following ZN-c3 treatment. **(I)** Representative flow cytometry plots of TCF1 versus Ly6C expression on PD-1+ CD8⁺ TILs from mice treated with the indicated doses of ZN-c3. **(J-K)** Quantification of PD-1⁺Ly6C⁺ (**J**), and TCF1⁺PD-1⁺ (**K**) populations among CD8⁺ TILs following ZN-c3 treatment. **(L–M)** Quantification of PD-1⁺Ly6C⁺ (**L**) and TCF1⁺PD-1⁺ (**M**) populations among CD4⁺ TILs following ZN-c3 treatment. **(N)** Experimental schematic illustrating combined ZN-c3 and FTY720 treatment in tumor-bearing mice. **(O)** Tumor growth curves for mEERL tumors in mice treated with vehicle or ZN-c3 (60 mg/kg) in the presence of FTY720. **(P)** Kaplan–Meier survival analysis of mEERL tumor–bearing mice treated with vehicle or ZN-c3 (60 mg/kg) in the presence of FTY720. Data are shown as mean ± SEM with individual data points representing biological replicates. Statistical significance was assessed using appropriate tests as indicated; *P < 0.05, **P < 0.01, ***P < 0.001; ns, not significant.

We next examined how WEE1i modulates immune cell populations *in vivo*. Because continuous high-intensity WEE1i induces tumor regression largely independent of adaptive immunity (**Fig. 2D-F**), whereas intermittent or submaximal dosing reveals immune-dependent mechanisms (**Fig. 2**; **Fig 3I-L**), we performed flow cytometric immunophenotyping of mEERL tumors treated with two submaximal ZN-c3 regimens (30 or 60 mg/kg) or vehicle control (**Fig. 5E**). ZN-c3 treatment increased overall immune cell infiltration relative to vehicle-treated tumors (**Fig. 5F**). Within the CD45^+^ leukocyte compartment, the abundance of CD3^+^ TILs increased at both dose levels (**Fig. 5G**). While FOXP3− (non-regulatory) CD4⁺ TILs expanded predominantly at the lower dose, CD8⁺ TILs and NK cells increased at both doses (**Fig. 5H**). CD8⁺ TILs exhibited an activated phenotype and lacked CD62L expression, and most expressed high levels of granzyme B, and upregulated inhibitory receptors including PD-1 and CD39 (**Sup. Fig. 5B**).

ZN-c3 treatment was associated with the expansion of two phenotypically distinct TIL populations within the TME, observed across both the CD8⁺ and CD4⁺ compartments. Within the PD-1⁺ gate, these populations were defined by a PD-1⁺TCF1⁺ subset and a PD-1⁺TCF1⁺Ly6C⁺ subset. In CD8⁺ TILs, both populations exhibited reduced expression of terminal exhaustion markers, including Eomes, Lag3, and Tim3, relative to vehicle-treated controls (**Fig. 5I–K** and **Sup. Fig. 5C**). A parallel expansion of PD-1⁺TCF1⁺ and PD-1⁺TCF1⁺Ly6C⁺ populations was also observed among CD4⁺ TILs (**Fig. 5L, M**).

PD-1⁺TCF1⁺ CD8⁺ and CD4⁺ T cells are known to arise in tumor-draining lymph nodes, where they function as a self-renewing progenitor reservoir that sustains intratumoral effector responses and is essential for durable tumor control during immunotherapy (*30–32*). To determine whether ZN-c3–induced antitumor immunity could be sustained by pre-existing TILs or instead required ongoing recruitment from secondary lymphoid organs, we pharmacologically blocked lymphocyte egress using FTY720. This intervention attenuated the therapeutic efficacy of ZN-c3, indicating that circulating T cells—and not solely resident TILs—contribute to WEE1i-mediated immune control (**Fig. 5N–P**). Consistent with these findings, ZN-c3 treatment did not increase the frequency of PD-1⁺TCF1⁺ TILs in human HPV⁺ OPC tumor slice cultures *ex vivo* (**Sup. Fig. 6F, G**). Together, these data indicate that WEE1i promotes infiltration and maintenance of cytotoxic lymphocytes with features of functional persistence in both murine and human HPV⁺ OPC tumors.

**Figure 6.**
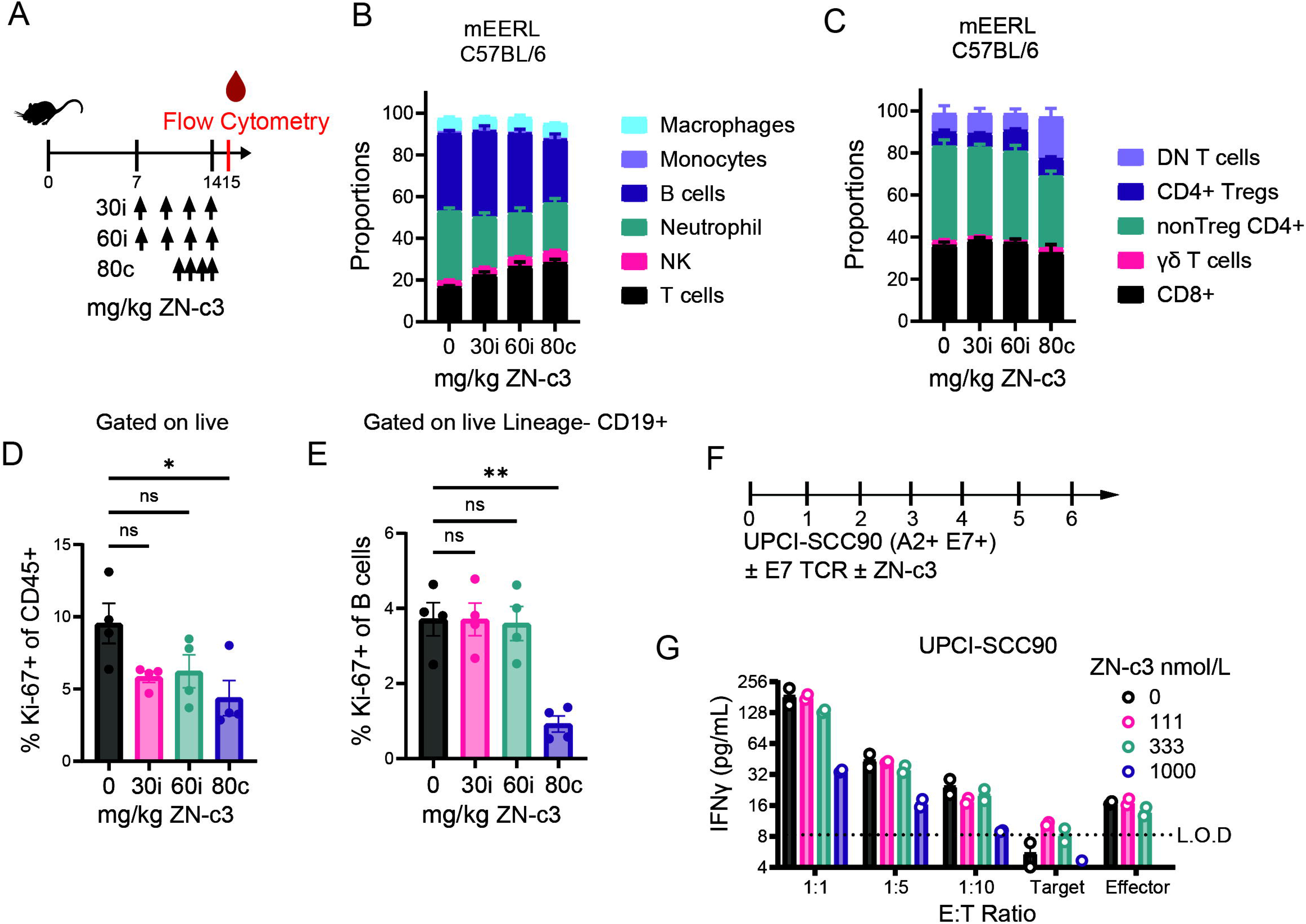
ZN-c3 dosing modulates systemic immune cell composition and proliferation. **(A)** Schematic of the *in vivo* ZN-c3 dosing regimen and experimental timeline. Tumor-bearing mice received ZN-c3 intermittently (i) or continuously (c) at the indicated doses, followed by peripheral blood collection and flow cytometric analysis. **(B)** Proportional composition of major immune cell populations among circulating leukocytes from mEERL tumor–bearing C57BL/6 mice treated with increasing doses of ZN-c3. **(C)** Proportional composition of circulating T cell subsets from the same animals. **(D)** Frequency of Ki-67⁺ cells among total CD45⁺ circulating leukocytes, gated on live cells, following treatment with the indicated doses of ZN-c3. **(E)** Frequency of Ki-67⁺ cells among circulating B cells, gated on live Lineage⁻ CD19⁺ cells, following treatment with the indicated doses of ZN-c3. **(F)** Experimental schematic for *in vitro* co-culture assays using UPCI-SCC90 (HLA-A2⁺, HPV16 E7⁺) target cells with E7-specific TCR-transduced T cells, treated with ZN-c3 as indicated. **(G)** IFN-γ concentrations in supernatants from **(F)** at the indicated effector-to-target (E:T) ratios and ZN-c3 concentrations. The dotted line denotes the limit of detection (L.O.D.). Data are shown as mean ± SEM with individual data points representing biological replicates. Statistical comparisons were performed using appropriate tests as indicated; *P < 0.05, **P < 0.01; ns, not significant.

### Effect of WEE1i dosing regimen on circulating immune cells

Our data indicate that ZN-c3 can mediate tumor control through distinct mechanisms depending on dosing schedule: continuous WEE1i achieves tumor regression largely independent of adaptive immunity, whereas intermittent dosing critically requires an intact immune system. We therefore asked whether these divergent dependencies are associated with differential effects on systemic immune cell populations. To address this, we evaluated the impact of intermittent ZN-c3 dosing (30 or 60 mg/kg) versus continuous dosing (80 mg/kg) on circulating immune cells in tumor-bearing mice (**Fig. 6A**).

ZN-c3 treatment induced a dose-dependent shift in circulating immune cell composition, characterized by an increased relative frequency of T cells that coincided with a reduction in neutrophils across all dosing regimens, including continuous dosing at 80 mg/kg (**Fig. 6B**). In contrast, reductions in B cell relative frequency were largely restricted to continuous dosing at 80 mg/kg. Within the T cell compartment, the relative frequencies of CD8⁺ T cells and non-regulatory CD4⁺ T cells were largely unchanged across dosing regimens, with the exception of continuous dosing at 80 mg/kg (**Fig. 6C**). This change was driven by an increased frequency of double-negative (CD4⁻CD8⁻) T cells observed only under continuous dosing at 80 mg/kg (**Fig. 6C**). No significant changes were observed in the relative frequencies of circulating monocytes, macrophages, regulatory T cells, or γδ T cells across dosing regimens (**Fig. 6B, C**).

To determine how dosing schedule influences immune cell proliferation and viability, we assessed Ki-67 expression and PARP cleavage (cPARP) across leukocyte subsets. Intermittent ZN-c3 treatment had minimal impact on immune cell proliferation, whereas continuous high-dose treatment (80 mg/kg) significantly reduced proliferation, most notably among B cells (**Fig. 6D, E**). This finding was consistent with the relative decrease in circulating B cell abundance observed at this dose (**Fig. 6B**). Similarly, continuous ZN-c3 treatment resulted in the greatest increase in cleaved PARP across immune cell populations, particularly in B and T cells (**Sup. Fig. 7A-D**). Together, these data indicate that intermittent WEE1i better preserves circulating immune cell composition compared with continuous high-dose dosing.

**Figure 7.**
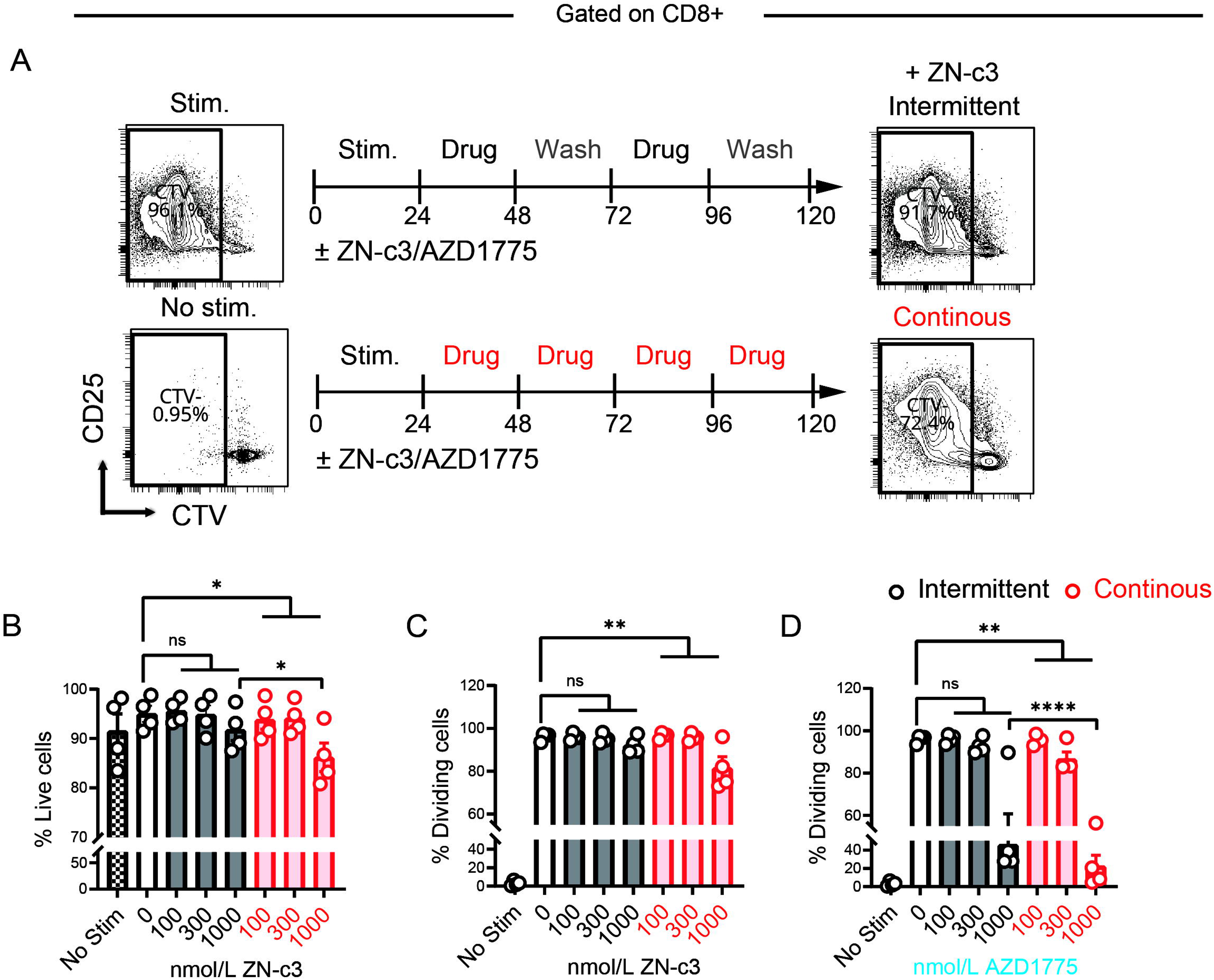
Intermittent WEE1i preserves T cell viability and proliferative capacity compared with continuous exposure. Human T cells in PBMC cultures were labeled with CTV and stimulated, exposed to ZN-c3 or AZD1775 for 5 days, and either maintained in drug (continuous) or subjected to drug washout (intermittent) prior to analysis by flow cytometry. **(A)** Schematic illustrating intermittent versus continuous *in vitro* exposure of stimulated human CD8⁺ T cells in PBMC cultures to the WEE1 inhibitors ZN-c3 or AZD1775. Representative flow cytometry plots show CTV dilution and CD25 expression at the experimental endpoint ± ZN-c3. **(B)** Quantification of CD8⁺ T cell viability following intermittent (gray) or continuous (red) exposure to increasing concentrations of ZN-c3. **(C)** Frequency of dividing CD8⁺ T cells following intermittent or continuous ZN-c3 exposure, assessed by CTV dilution. **(D)** Frequency of dividing CD8⁺ T cells following intermittent or continuous exposure to AZD1775. Data are representative of ≥3 independent experiments with individual data points shown. Bars represent mean ± SEM. Statistical significance was determined using appropriate multiple-comparison tests; *, P < 0.05, **, P < 0.01, ****, P < 0.0001; ns, not significant.

### Intermittent ZN-c3 dosing preserves T cell proliferation and function

To directly assess how ZN-c3 schedule influences T cell function, we cocultured human E7 TCR–transduced CD8⁺ T cells with UPCI-SCC90 cells (HLA-A*02:01⁺ HPV⁺ OPC) across a range of effector-to-target ratios in the presence of increasing concentrations of ZN-c3 for 6 days. Consistent with our *in vivo* observations, sustained ZN-c3 exposure resulted in dose-dependent attenuation of T cell effector activity, as measured by IFN-γ secretion (**Fig. 6F, G**). These findings indicate that prolonged WEE1i can constrain T cell function and suggest that limiting exposure duration may mitigate these effects.

To more precisely define how dosing schedule influences T cell proliferation and signaling, human peripheral blood mononuclear cells (PBMCs) were activated with anti-CD3 and anti-CD28 antibodies and treated with ZN-c3 under continuous or intermittent regimens. ZN-c3 induced a dose-dependent reduction in inhibitory CDK1 phosphorylation, accompanied at the highest concentration (1,000 nmol/L) by accumulation of γH2AX. Both pCDK1 suppression and γH2AX accumulation were fully reversible following drug washout (**Sup. Fig. 8A–C**), indicating transient cell cycle checkpoint engagement rather than irreversible T cell injury.

Using CellTrace Violet (CTV) dilution assays, we quantified T cell proliferation as the percentage of cells undergoing division. Continuous ZN-c3 exposure at 1,000 nmol/L was associated with a modest but statistically significant reduction in the fraction of dividing CD4⁺ and CD8⁺ T cells, whereas no significant changes in proliferative fraction were observed under intermittent dosing at the same concentration (**Fig. 7A, C; Sup. Fig. 8D**). The reduction in dividing cells under continuous exposure was limited in magnitude (approximately 20%) and was accompanied by only a modest decrease in T cell viability (approximately 10–15%), indicating largely preserved T cell survival (**Fig. 7B; Sup. Fig. 8F**). In parallel experiments, ZN-c3 effectively suppressed proliferation of HPV⁺ OPC cell lines at similar or lower concentrations, supporting a dosing window in which intermittent WEE1i preferentially preserves T cell proliferative capacity relative to tumor cells (**Sup. Fig. 9A**).

Consistent with these observations, the first-generation WEE1 inhibitor AZD1775 suppressed T cell proliferation more profoundly than ZN-c3 at comparable doses, indicating a narrower immune tolerability profile (**Fig. 7D; compare with Fig. 7C**). Analysis of dividing T cells confirmed that intermittent dosing was better tolerated than continuous exposure for both inhibitors, with ZN-c3 exhibiting a more favorable profile overall (**Fig. 7C, D; Sup. Fig. 8D, E**). Phenotypic analysis of cycling (CTV low) CD8⁺ T cells revealed that granzyme B expression remained unchanged within the dividing population, while markers associated with T cell exhaustion (TOX, TIM-3, LAG-3) and activation (CD25) were similarly maintained (**Sup. Fig. 8G**). These data indicate that reduced proliferation under continuous WEE1 inhibition occurs without induction of canonical exhaustion programs, impaired activation, or loss of cytotoxic effector differentiation among responding T cells.

Collectively, these findings indicate that continuous WEE1i imposes a cumulative, yet reversible, constraint on T cell expansion, whereas intermittent ZN-c3 dosing preserves proliferative competence and cytotoxic effector features across both CD4⁺ and CD8⁺ T cell compartments, providing a mechanistic rationale for improved immune tolerability under intermittent dosing schedules.

## Discussion

We found that WEE1i anti-HPV⁺ OPC activity involves two distinct mechanisms: 1) tumor cell–autonomous cytotoxicity, and 2) modulation of antitumor immunity. HPV⁺ tumor cells are highly sensitive to WEE1i, in part because they rely upon the G2/M checkpoint to tolerate oncogene-driven genomic instability, which is disabled by inhibiting WEE1 (*6–10*). Our previous work demonstrated the role of the FOXM1-CDK1 axis in driving WEE1i-sensetivity in HPV+ models of OPC (*9*), as well as early clinical promise of an AZD1775-based regimen in advanced HPV+ OPC (*10*). Here, we extend these findings by demonstrating a critical role for host immunity in the antitumor activity of both AZD1775 and the next-generation WEE1 inhibitor ZN-c3, which achieved durable responses as single-agents in immunocompetent HPV⁺ OPC models (**Fig. 1, 2**). Comparison of these two structurally distinct WEE1 inhibitors revealed shared antitumor and immune phenotypes consistent with on-target WEE1 inhibition, as well as differences in immune tolerance between compounds.

WEE1i elicits durable antitumor immunity that depends on CD4⁺ and CD8⁺ T cells and cDC1s, implicating antigen-specific immune responses in therapeutic efficacy (**Fig. 2**). HPV⁺ OPC is an immunogenic malignancy, attributable to expression of viral antigens and other mechanisms (*33*). While current HPV-targeted T cell therapies largely focus on E6 and E7 (*34*), our findings suggest that the immunologic effects of WEE1i extend beyond simply targeting viral epitopes, and also involves complex interplay between DNA damage, innate immune sensing (cGAS/STING), and cDC1-dependent priming within a permissive TME (**Fig. 2-4**). WEE1i may thus increase antigen availability and broaden antitumor T cell specificity. Future studies incorporating additional HPV antigens or full viral genome expression will be important to define the breadth of immunity elicited by WEE1i.

In contrast to reports in other solid tumor models (*11–13*), WEE1i does not directly activate type I IFN signaling within HPV⁺ tumor cells (**Fig 3E, F**), but instead provokes a type I IFN response within STING-proficient non-tumor cells in the TME (**Fig. 3G-J**; **Fig. 4H**). This mechanism may be unique to HPV⁺ cancers, as HPV oncoproteins actively suppress STING signaling, inhibit IRF3 activation, and dampen type I IFN production within infected epithelial cells (*15*). Accordingly, we find that HPV⁺ tumor cells are intrinsically STING-deficient, yet intact STING signaling within the host compartment is essential for the antitumor immunity induced by WEE1i (**Fig. 3E-J**).

While host STING signaling is required, tumor cell–intrinsic cGAS remains necessary for therapeutic efficacy (**Fig. 3K, L**). WEE1i induces DNA damage and activate cGAS–STING signaling in cDC1s within solid tumor models (*12*), providing a potential explanation for IFN responses we observe following ZN-c3 treatment *ex vivo* (**Fig. 3G, H**). Indeed, despite the potential engagement of innate sensing pathways in myeloid cells, we find that cGAS deletion within tumor cells ablates productive antitumor immune responses following WEE1i (**Fig. 3K, L; Sup. Fig. 3I, J**). These findings support a hierarchical model in which WEE1i-induced DNA damage is initially sensed within tumor cells and subsequently amplified by STING-competent host cells within the TME (**Fig. 4H**). Such tumor-to-host cGAS–STING communication has been described in other contexts of DNA damage–induced immunity (*35–37*).

Overall, WEE1i reshapes the HPV⁺ OPC TME to favor cytotoxic lymphocyte recruitment, function, and persistence (**Fig. 4 and 5**). The antitumor efficacy of WEE1i in our preclinical models was associated with enhanced lymphocyte-mediated immune activity. A key finding of this study is that the WEE1 inhibitor ZN-c3 is both immune-permissive and immune-modulatory, defining a favorable preclinical safety profile. Unlike continuous WEE1i, which impairs T cell proliferation and function (*11, 18*), intermittent dosing preserves T cell fitness while maintaining antitumor efficacy (**Fig. 1 and 7**). This distinction is critical, as T cells are themselves highly proliferative and vulnerable to perturbations in cell-cycle regulation (*38*).

The increase in PD-1⁺TCF1⁺ CD8⁺ and CD4⁺ TILs (**Fig 5. I, K, M**) aligns with the concept that progenitor-like cells can be mobilized from secondary lymphoid organs to sustain TME immunity (*31, 32*), which may help explain the sensitivity of intermittent dosing to lymphocyte trafficking (**Fig. 5N-P**). In parallel, intermittent WEE1i was associated with the emergence of PD-1⁺Ly6C⁺ CD8⁺ TILs that retained TCF1 expression (**Fig. 5I, J**), consistent with an activated yet progenitor-like CD8 state. Consistent with recent reports in other solid tumors (*39–41*), the accumulation of Ly6C-expressing CD8⁺ TILs has been associated with enhanced antitumor immunity and improved tumor control, potentially reflecting IFN-driven programs that support lymphoid homing, central memory differentiation, and preservation of proliferative capacity (*40, 42, 43*). Together, these features may contribute to durable immune surveillance in HPV⁺ OPC. This is particularly relevant in HPV⁺ OPC, where tumors harbor antigen-specific CD8⁺ T cells recognizing a broad repertoire of viral antigens, including E2 and E5, and spanning diverse differentiation states with variable proliferative capacity (*30*). Preservation of T cell proliferative potential during therapy may therefore be a critical determinant of durable immune-mediated tumor control in this setting.

Patients with recurrent unresectable or metastatic HPV⁺ OPC have incurable disease and limited therapeutic options. Induction immune checkpoint blockade (ICB) prior to definitive treatment is a emerging strategy for recurrence-prone HPV⁺ OPC, with recent phase II trials demonstrating proof-of-concept but also highlighting the need for better combination approaches (*44, 45*). FDA-approved immune checkpoint inhibitors have suboptimal response rates in both HPV-related and HPV-unrelated head and neck cancers (*46*), indicating that additional strategies are needed to further augment immunotherapeutic benefit. As an orally bioavailable agent with a favorable systemic toxicity profile and emerging immunomodulatory activity, WEE1i represents a compelling therapeutic approach for HPV⁺ OPC. Our findings demonstrate that WEE1 inhibitors can function not only as direct antitumor agents but also as immune-enabling therapies, providing a rationale for combination strategies with immune checkpoint blockade or HPV-directed T cell–based therapies to improve outcomes in advanced disease. The therapeutic potential of this agent may also extend to the curative intent setting where toxicity mitigation and treatment de-escalation for low-risk disease is an active area of clinical investigation.

## Methods

### Compound Preparation and Treatment

ZN-c3 (Zentalis Pharmaceuticals, Inc.) and AZD1775 (Selleck) were formulated in 0.5% methylcellulose and administered by oral gavage as indicated in the figure legends.

### Cell Lines

mTE, mEERL, mEE-F3T3 and MLM3 cells were maintained in a 1:3 mixture of Ham’s F-12 and DMEM supplemented with 10% fetal bovine serum (FBS), 1% penicillin/streptomycin, 50 μg/mL hydrocortisone, 5 μg/mL insulin–transferrin–selenium, 0.65 ng/mL triiodothyronine, and 5 ng/mL EGF (E-media). Human cell lines were cultured in DMEM supplemented with 10% FBS and 1% penicillin/streptomycin (Supplementary Table 1). All cell lines were maintained at 37°C in a humidified atmosphere with 5% CO₂ and periodically tested and confirmed negative for Mycoplasma.

### Animal Studies

WT C57BL/6J, Batf3⁻/⁻, Sting^gt/gt^ and NOD/SCID/IL2Rγ⁻/⁻ mice were purchased from the Jackson Laboratory and/or bred at the University of Pennsylvania and housed under specific pathogen-free conditions at the University of Pennsylvania and the Fred Hutchinson Cancer Center. For tumor implantation studies, mice aged 8–12 weeks were used. Experiments were randomized but not blinded. All animal experiments were conducted in accordance with institutional guidelines and were approved by the Institutional Animal Care and Use Committees of the University of Pennsylvania (IACUC #807486) and the Fred Hutchinson Cancer Center (IACUC #1413).

For subcutaneous flank tumors, mice received 1 × 10⁶ tumor cells in 100 μL PBS/Matrigel (2:1) subcutaneously in the right flank. For orthotopic buccal mucosa model, mice were anesthetized with isoflurane, and 0.5 × 10⁶ cells in 50 μL of PBS/Matrigel (2:1) were injected submucosally into the left buccal mucosa. Tumor dimensions were measured using digital calipers, and volume was calculated as (length × width²)/2. Mice were euthanized per institutional humane endpoints. For rechallenge studies, mice with established orthotopic mEERL tumors that achieved CR following ZN-c3 monotherapy and remained tumor-free for at least 90 days were rechallenged with 1 × 10⁶ mEERL cells implanted in the contralateral flank. Age- and sex-matched, tumor-naïve WT B6 mice served as controls.

### Generation of cGAS-Knockout Cell Lines

cGas knockout mEERL cells were generated using CRISPR–Cas9–mediated genome editing with the pSCAR system (*47*). Lentiviral particles encoding Cas9-GFP (pSCAR_Cas9; Addgene #162074) were produced in 293T cells using psPAX2 (#12260) and pMD2.G (#12259), both gifts from Didier Trono, and used to transduce mEERL cells. FACS-enriched GFP⁺ Cas9-expressing mEERL cells were subsequently transfected with a pooled set of three synthetic single guide RNAs (sgRNAs) targeting distinct regions of murine *cGas* (Supplementary Table 2) or transduced with pSCAR_sgcGAS lentivirus, and edited cells were enriched by FACS (mKate⁺). Cre-mediated excision was subsequently performed using integrase-deficient lentivirus expressing Cre recombinase (psPAX2-D64V; Addgene #63586) to generate GFP⁻mKate⁻ knockout cells. Single cell clones were obtained by limiting dilution and knockout was confirmed by genomic sequencing and immunoblotting. For *in vivo* studies, 1 × 10⁶ cGAS-KO cells were subcutaneously implanted into WT B6 mice.

### Tumor Slice Cultures

For murine tumor slice cultures, mEERL tumors (∼1,000mm³) were embedded in 4% low-melt agarose and sectioned at 250-300 μm using a Compresstome (VF-510-0Z; speed 6, oscillations 4). Slices were cultured on cell culture inserts (Millicel, Millipore) in 24-well plates in E-media at 37°C/5% CO₂.

For human tumor slice cultures, de-identified treatment-naïve HPV^+^ oropharyngeal squamous cell carcinoma specimens were obtained from patients undergoing surgical resection under an IRB-approved protocol (protocol No. 417200) with written informed consent. Age, sex and specimen descriptions are in Supplementary Table 3. Fresh specimens were processed within 2 hours and sectioned as above.

### Flow Cytometry Analysis

Samples were processed in a 96-well plate format. Non-spleen samples were concentrated to 200 μL via centrifugation and vacuum aspiration, while spleen samples were directly resuspended to 200 μL. Cells were stained with Zombie NIR™ dye or LIVE/DEAD™ Fixable Blue Dead Cell Stain and CD16/CD32 blocking antibody for 10 minutes at room temperature (RT). Surface staining was performed in True-Stain™ Multi-Fluor or BD Horizon™ Brilliant Stain Buffer for 30-60 minutes at RT.

After surface staining, cells were fixed with fixation buffer (eBioscience) for 30 minutes at RT and washed with Permeabilization buffer. For intracellular staining, cells were fixed and permeabilized using Foxp3/Transcription Factor Staining Buffer Set (eBioscience) per the manufacturer’s guidelines and subsequently stained for 1 hour at RT or overnight at 4°C. Cells were then washed with Permeabilization buffer and resuspended in 200 μL FACS buffer (PBS + 2% FBS + 0.04% EDTA) for analysis. Single-color compensation controls were prepared by labeling one drop of UltraComp eBeads™ Compensation Beads with 0.5–1 µL of each antibody (Supplementary Table 4-6). Data were acquired on a Cytek Aurora or LSR Fortessa flow cytometer and analyzed using OmiQ or FlowJo version 10.10.0.

### In Vitro T-Cell Functional Assays

PBMCs and purified CD8^+^ T cells were obtained from healthy volunteers with informed consent obtained from all subjects in this study and apheresis was collected under University of Pennsylvania IRB-approved protocol 705906 and processed by the Human Immunology Core (HIC). PBMCs were labeled with CellTrace Violet (5 μmol/L) and were activated and expanded using the bead-free ImmunoCult™ Human CD3/CD28/CD2 T Cell Activator (STEMCELL Technologies) or anti-CD3/CD28 microbeads (Life Technologies). Cells were treated with ZN-c3 or AZD1775 as indicated, and proliferation was assessed by dye dilution using flow cytometry.

### Single-Cell RNA Sequencing

mEERL tumors from B6 mice treated with AZD1775 (60 mg/kg, 4 doses over 8 days) or vehicle were processed using the Tissue Fixation and Dissociation for Chromium Fixed RNA Profiling protocol (CG000553, 10X Genomics) and the Chromium Next GEM Single Cell Fixed RNA Sample Preparation Kit (1000414, 10X Genomics). Libraries were sequenced on an Illumina NextSeq2000 platform for 15,000 reads per cell. Raw sequencing data was processed using 10X Genomics CellRanger with mm10 (GRCm38) as reference genome for specified panel of 17,348 genes and then postprocessed in R using Seurat v4. Cells with fewer than 500 reads and mitochondrial gene percentage of >20% were filtered out. Filtered cells were normalized, scaled and clustered in Seurat v4. Cell type annotation was performed by SingleR using the mouse-cell specific markers established in ImmGen Project. Gene set enrichment analysis (GSEA) was performed at pseudobulk level to assess pathway-level changes across compartments (immune, stromal, tumor).

### Cell-Cell Interaction Analysis

Ligand-receptor interactions were inferred from scRNA-seq data using CellChat v1.6.1 (*27*). Signaling networks were compared between treatment groups, focusing on antigen-presenting cell populations (dendritic cells, tumor-associated macrophages, tumor-associated neutrophils) as senders and lymphocyte populations (CD4^+^ T cells, CD8^+^ T cells, NK cells) as receivers. Pathway-specific communication probabilities were calculated and visualized.

### STING Pathway Functional Assays

mEERL and mTE cells were treated with the STING agonist diABZI (10 μmol/L, InvivoGen) or transfected with poly(I:C) (1 μg/mL, InvivoGen) or double-stranded DNA analog (ISD Naked, InvivoGen) using polyethylenimine for 48 hours. Gene expression was assessed by RT-qPCR, and STAT1 phosphorylation was measured by western blot as described above.

### T-Cell Depletion and FTY720 treatment

CD4^+^ and CD8^+^ T cells were depleted using intraperitoneal injections of depleting antibodies: anti-CD4 (clone GK1.5, Bio X Cell, 200 μg), anti-CD8α (clone 2.43, Bio X Cell, 200 μg), or combined anti-CD4/CD8 for pan-T-cell depletion. Antibodies were administered 1 day before ZN-c3 initiation and twice weekly thereafter. For FTY720 experiment, FTY720 (MedChemExpress) was provided in the drinking water (2 µg/mL) on day 7 post inoculation. FTY720 treatment was continued throughout the entire experimental course.

### Lentiviral TCR construct

TCR HPV E7 (Addgene #122728) lentiviral construct was generated using the third-generation lentiviral transfer vector pTRPE-eGFP-T2A-mCherry (provided by Michael C. Milone, University of Pennsylvania).

### TCR HPV E7-engineered T cells

Purified human CD8^+^ T cells were obtained from healthy volunteers through HIC as detailed above. TCR HPV E7-engineered CD8+ T cells were generated as described previously (*48*). Briefly, primary CD8+ T cells were activated with anti-CD3/CD28 microbeads (Life Technologies) and transduced with TCR E7 lentiviral vector particles at an MOI of 5. Cell cultures were grown in IL-7 (5 ng/ml; R&D Systems) / IL-15 (5 ng/ml; R&D Systems) supplemented T cell media. On day 5, microbeads were removed and T cells electroporation with sgRNAs targeting *TRAC* and *TRBC1/TRBC2* (Supplementary Table 2, Integrated DNA Technologies) complexed with TrueCut Cas9 Protein V2 (10 µg, ThermoFisher Scientific). Edited T cells were cultured in IL-7/IL-15 supplemented T cell media for a total of 14 days followed by Rapid Expansion Method (*49*). HPV16 E7-specific T cell frequencies were determined by p-HLA multimer flow cytometric assays.

### In vitro cytotoxicity Assay

For cytotoxicity assays, E7 TCR-transduced CD8+ T cells were cocultured with UPCI-SCC90 target cells at effector: target ratios of 1:1 to 1:10 in the presence of ZN-c3 for 6 days. Culture supernatants were collected for IFN-γ quantification by ELISA (Human IFN-γ DuoSet, R&D Systems).

### Adoptive T-Cell Therapy

LNT-20 PDXs were established and passaged in the subcutaneous flank of NSG mice as described (*50*). Tumor volumes were measured as above. Treatment started when tumors reached ∼150-250 mm^3^. Mice bearing established LNT-20 tumors received intravenous injection of 0.5-4 × 10⁶ E7 TCR T cells intravenously with or without ZN-c3 treatment.

## Statistical Analysis

Data are presented as mean ± SEM unless otherwise indicated. Statistical analyses and graphical representations were performed using GraphPad Prism (RRID: SCR_002798) version 9 or 10 and R (v4.2). Tumor growth significance was assessed using growth-rate–based modeling of longitudinal tumor volume measurements, fitting individual tumors to an exponential growth model and comparing estimated growth rates between treatment groups as described previously (*51*). Survival analyses were performed using Kaplan–Meier curves with log-rank (Mantel–Cox) tests.

Flow cytometry data, gene expression measurements, and other continuous variables were analyzed using unpaired two-tailed Student’s *t* tests (for two-group comparisons) or one-way ANOVA with appropriate post-hoc multiple-comparisons tests (for three or more groups), as indicated in the figure legends. When assumptions of normality were not met, nonparametric tests were used as noted. A *P* value < 0.05 was considered statistically significant. Sample sizes were determined based on prior studies and institutional guidelines; no samples were excluded from analysis.

## Acknowledgement

We thank Anna Elz and the Fred Hutch Innovation lab for technical help with the scRNA-seq experiment. We thank the CHOP Flow Cytometry Core Laboratory (RRID:SCR_009726) for technical expertise and access to instrumentation. We also acknowledge the University of Pennsylvania Perelman School of Medicine Molecular Pathology and Imaging Core (MPIC; RRID:SCR_022420), the Stem Cell and Xenograft Core (RRID:SCR_010035), and HIC (RRID:SCR_022380) for their support. This research was also supported by NIH P30 CA015704 from the Fred Hutchinson Cancer Center/University of Washington/Seattle Children’s Cancer Consortium, which supports the Experimental Histopathology Shared Resource (RRID:SCR_022612), the Comparative Medicine Shared Resource (RRID:SCR_022610), and the Genomics and Bioinformatics Shared Resource (RRID:SCR_022606).

This work was supported by a Stand Up To Cancer (SU2C)–Catalyst Award (to B.E.C., D.B., and A.D.). SU2C is a division of the Entertainment Industry Foundation, and research grants are administered by the American Association for Cancer Research, the Scientific Partner of SU2C.

Additional funding was provided by the Fred Hutch Pathogen-Associated Malignancies Integrated Research Center (PAM-IRC) Pilot Award (to A.D.). A.D. is supported by the National Institutes of Health Pathway to Independence Award (K99/R00DE030194), and the American Cancer Society Institutional Research Grant IRG-22-150-41-IRG. D.B. was supported by NIH award R01DE034056, and the Stephen and Susan Kelly Family Fund for Head and Neck Cancers. A.D. and D.B. were supported by the Breakthrough Challenge Foundation, and the William and Greta Lydecker Fund.

## Data Availability Statement

The data generated in this study are available upon request from the corresponding author.

---

**Supplementary Figure 1.**
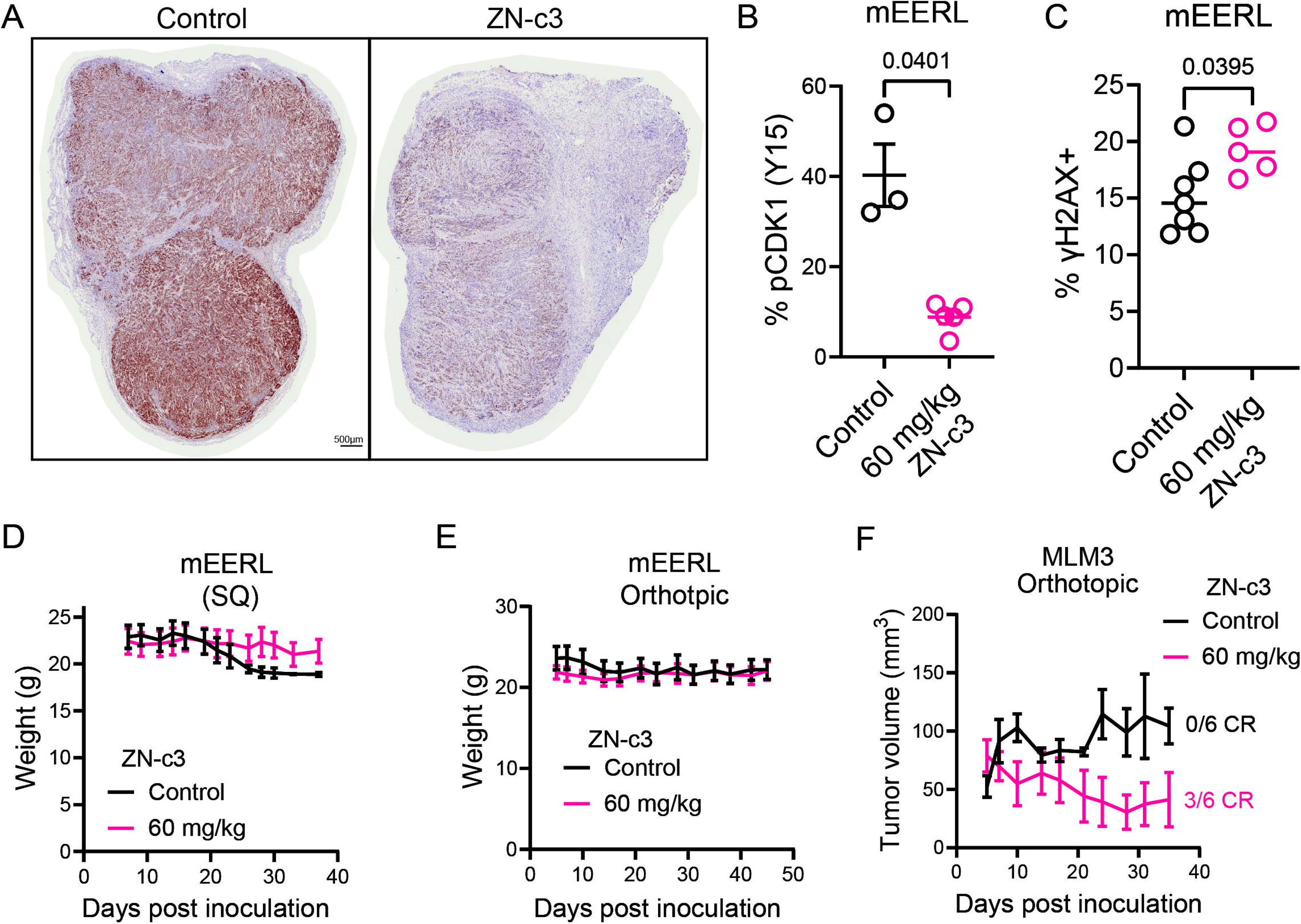
Effects of ZN-c3 on CDK1 phosphorylation, DNA damage, tumor growth, and body weight *in vivo*. (A) Representative immunohistochemical staining of phosphorylated CDK1 (pCDK1^Y15^) in mEERL tumors from mice treated with vehicle (Control) or ZN-c3 (60 mg/kg). (B) Quantification of pCDK1^Y15+^ cells in mEERL tumors from control and ZN-c3–treated mice. (C) Quantification of γH2AX^S139+^ cells in mEERL tumors following treatment with vehicle or ZN-c3 (60 mg/kg). Body weight curves of mice bearing subcutaneous (D) or (E) orthotopic mEERL tumors treated with vehicle or ZN-c3 (60 mg/kg). (F) Tumor growth curves of orthotopic MLM3 tumors from mice treated with vehicle or ZN-c3 (60 mg/kg). Data are presented as mean ± SEM where applicable. Statistical significance is indicated as shown.

**Supplementary Figure 2.**
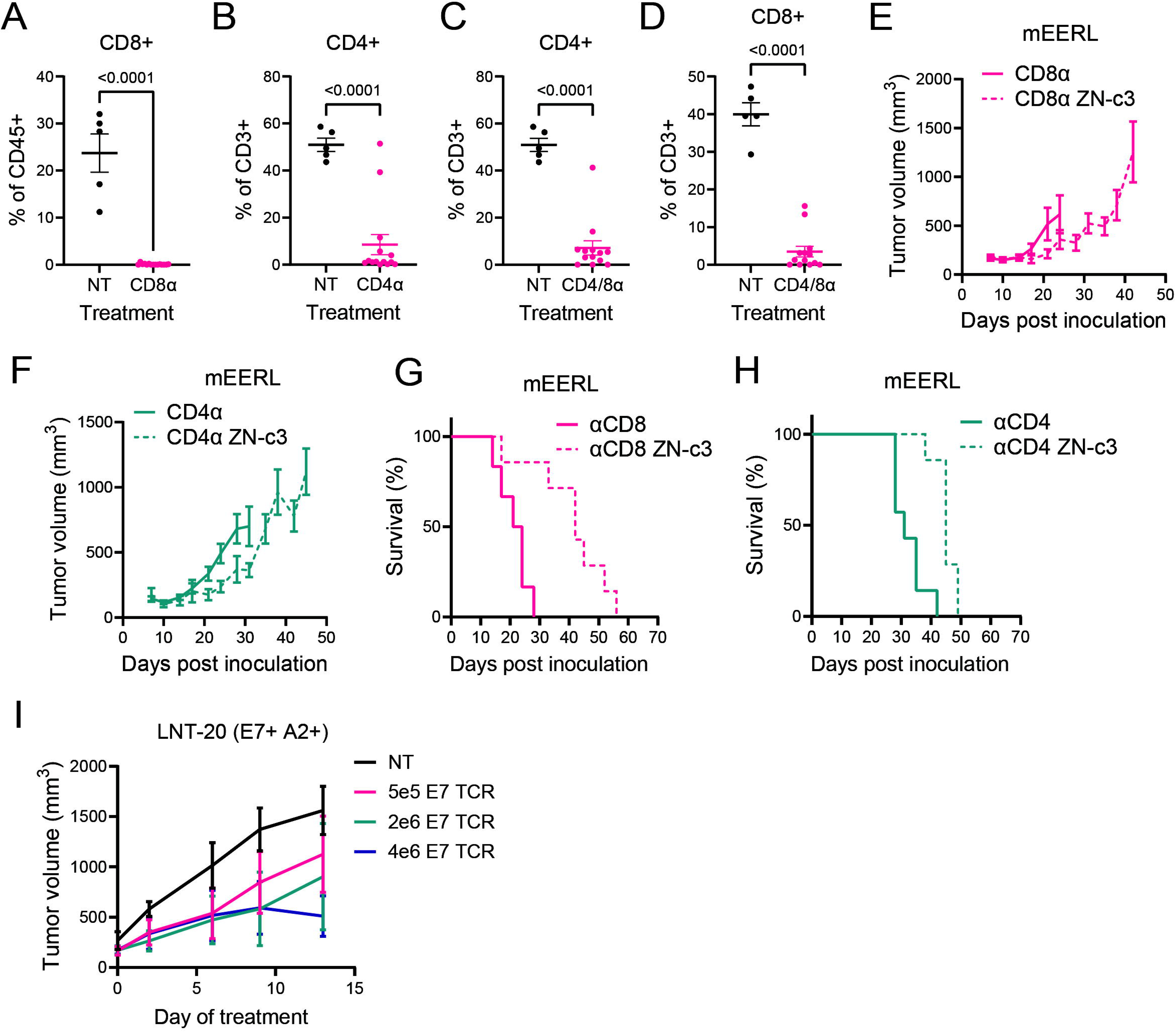
T cell depletion efficiency and tumor growth analyses during WEE1i in vivo. (A–D) Flow cytometric validation of *in vivo* T cell depletion. Frequencies of CD8⁺ T cells (A) and CD4⁺ T cells (B–D) among tumor-infiltrating leukocytes following treatment with depleting antibodies against CD8α or CD4α, compared to non-treated (NT) controls. Cells were gated on live CD45⁺ or CD3⁺ populations as indicated. Tumor growth curves of mEERL tumors in WT mice treated with anti-CD8 antibodies (E) or anti-CD4 antibodies (F) in the presence or absence of ZN-c3 treatment. Kaplan–Meier survival analysis of mEERL tumor-bearing mice treated with anti-CD8 antibodies (G) or anti-CD4 antibodies (H) in the presence or absence of ZN-c3 treatment. (I) Tumor growth curves of HPV16 E7⁺ HLA-A*02:01⁺ LNT-20 xenografts in NSG mice following adoptive transfer of increasing numbers of HPV16 E7–specific TCR-engineered T cells.

**Supplementary Figure 3.**
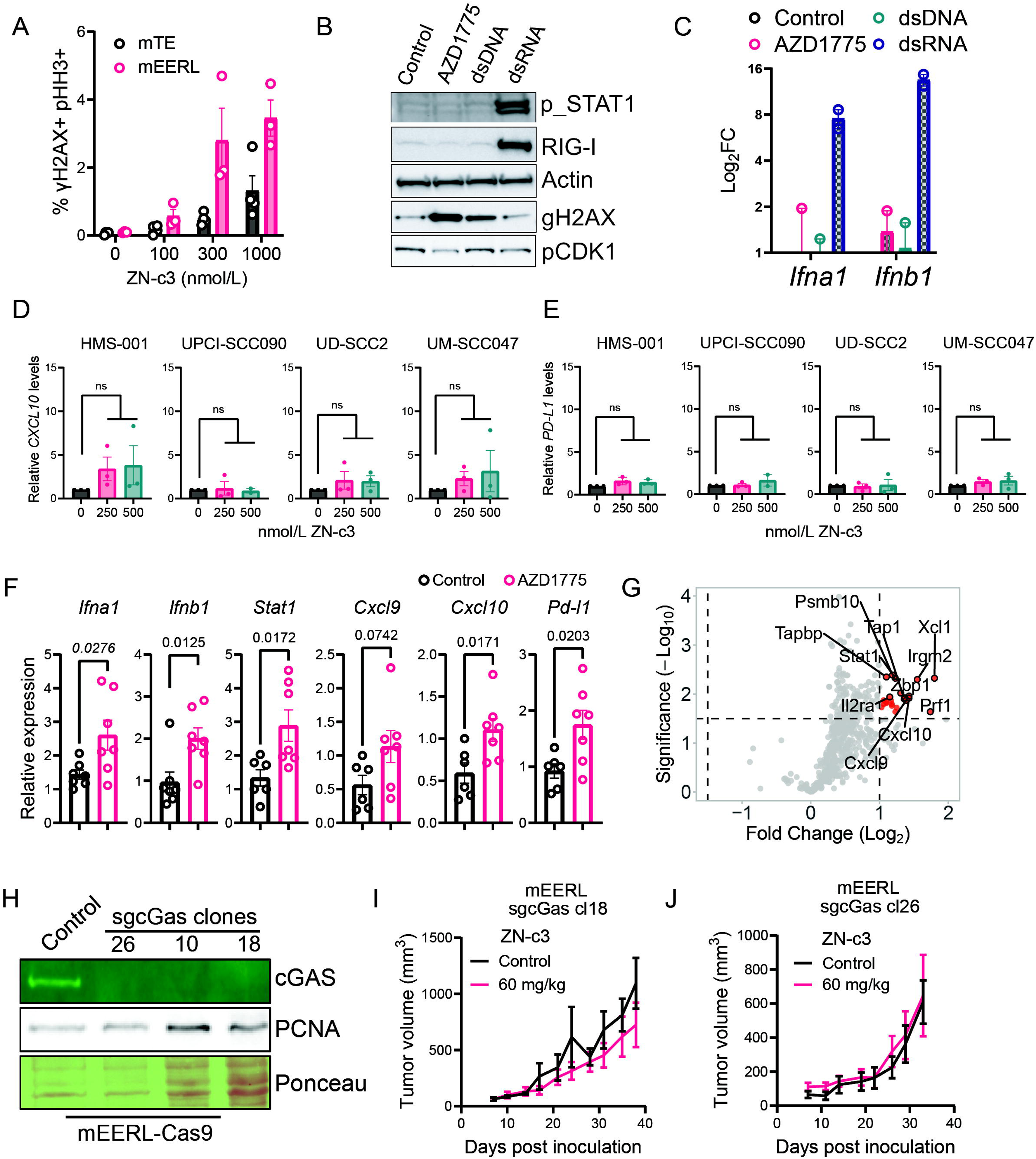
DNA damage markers and interferon-related gene expression following WEE1i *in vitro* and *in vivo*. (A) Flow cytometric quantification of γH2AX⁺ pHH3⁺ cells in isogenic mTE and mEERL cells treated *in vitro* with increasing concentrations of ZN-c3 for 24 hours. (B) Immunoblot analysis of pSTAT1, RIG-I, γH2AX, and pCDK1^Y15^ levels in mEERL cells treated with 250 nmol/L AZD1775 (WEE1i), transfected double-stranded DNA (dsDNA), or double-stranded RNA (dsRNA). (C) Relative mRNA expression (log₂ fold change) of *Ifna1* and *Ifnb1* in mEERL cells treated with AZD1775, dsDNA, or dsRNA compared to control. Relative *CXCL10* (D) and *CD274* (E) mRNA expression in HPV⁺ human head and neck cancer cell lines (HMS-001, UPCI-SCC090, UD-SCC2, UM-SCC047) treated with ZN-c3 (250 or 500 nmol/L). (F) Relative mRNA expression of interferon-stimulated and immune-related genes (*Ifna1, Ifnb1, Stat1, Cxcl9, Cxcl10,* and *Cd274*) in mEERL tumors following treatment with 60 mg/kg AZD1775 compared to control (four doses, 3x weekly). (G) Volcano plot of differentially expressed genes in mEERL tumors following AZD1775 treatment based on NanoString Profiling. (H) Immunoblot validation of cGAS protein loss in CRISPR-edited mEERL sg_cGas clones (clones 26, 10, and 18). (I, J) Tumor growth curves of mice bearing mEERL sg_cGas clone 18 (I) or clone 26 (J) tumors treated with vehicle or ZN-c3 (60 mg/kg, 3x weekly). Data are shown as mean ± SEM where applicable. Statistical significance was determined using unpaired t-test and is indicated as shown.

**Supplementary Figure 4.**
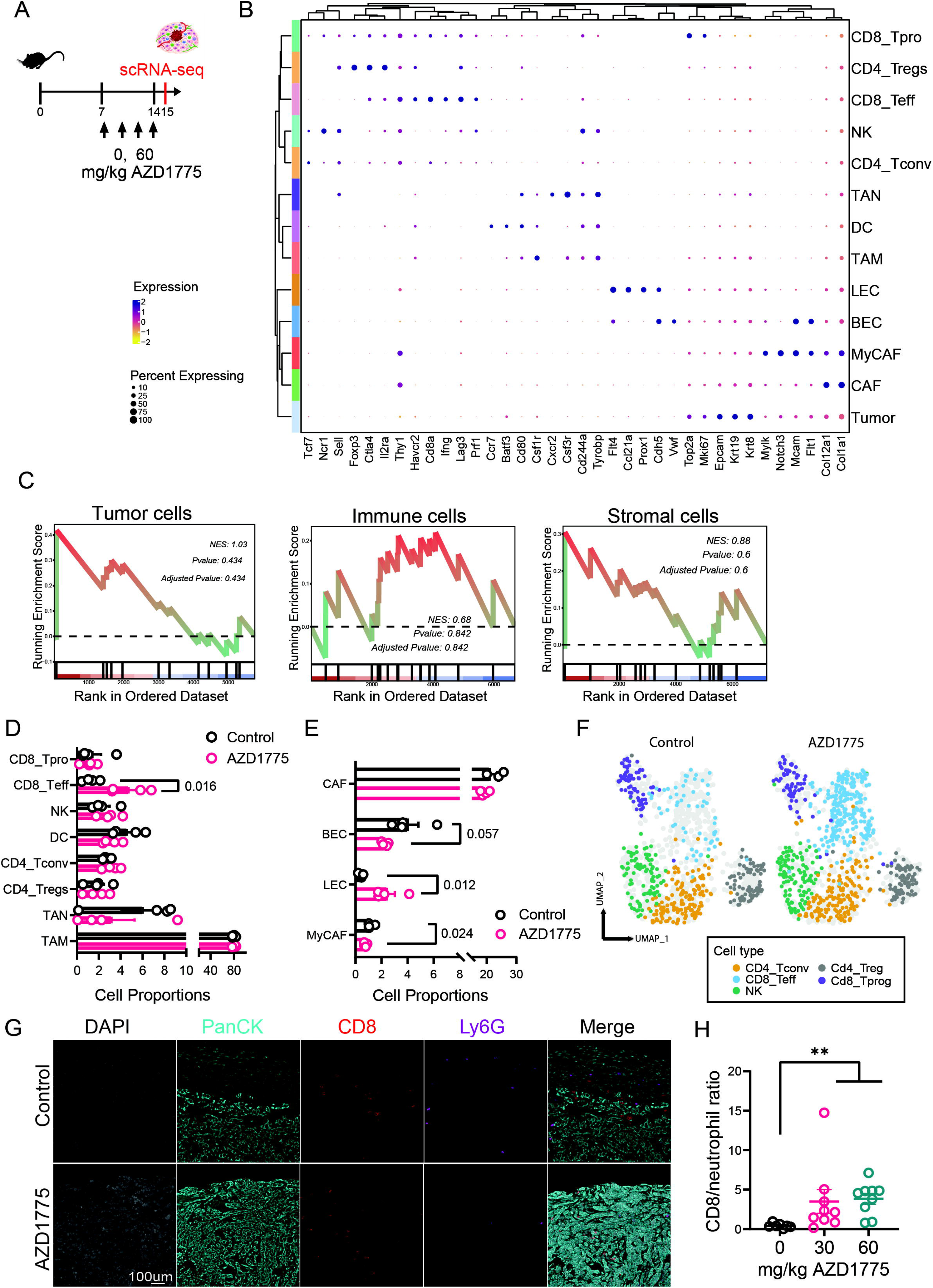
Extended immunophenotypic analysis of tumors following AZD1775 treatment. (A) Experimental schematic of the *in vivo* AZD1775 dosing regimen and timeline for scRNA-seq analysis of tumor-infiltrating immune cells. Tumor-bearing mice received AZD1775, or vehicle as indicated prior to tumor harvest. (B) Dot plot showing expression of canonical marker genes across major tumor, immune, and stromal cell populations identified by single-cell RNA sequencing of mEERL tumors. Dot size represents the percentage of cells expressing each gene, and color intensity indicates scaled average expression. (C) Gene set enrichment analysis (GSEA) plots showing integrated stress response hallmark pathway enrichment scores in tumor, stromal, and immune compartments following AZD1775 treatment, based on compartment-specific differential gene expression. Normalized enrichment scores (NES), nominal P values, and adjusted P values are shown. (D) Proportions of immune cell subsets identified by single-cell RNA sequencing in control and AZD1775-treated mEERL tumors, including CD8⁺ progenitor-like T cells (CD8_Tprog), CD8⁺ effector T cells (CD8_Teff), natural killer (NK) cells, dendritic cells (DCs), conventional CD4⁺ T cells (CD4_Tconv), and regulatory T cells (CD4_Tregs). (E) Proportions of stromal cell populations identified by single-cell RNA sequencing in control and AZD1775-treated tumors, including cancer-associated fibroblasts (CAF), blood endothelial cells (BEC), lymphatic endothelial cells (LEC), and myofibroblastic CAFs (MyCAF). (F) Uniform Manifold Approximation and Projection (UMAP) visualization of lymphocyte populations from control and AZD1775-treated tumors, colored by annotated cell type. (G) Representative multiplex immunofluorescence images of mEERL tumor sections from control and AZD1775-treated mice stained for DAPI (nuclei), Pan-cytokeratin (PanCK; tumor epithelium), CD8 (T cells), and Ly6G (neutrophils). Scale bar, 100 μm. (H) Quantification of the intratumoral CD8⁺ T cell–to–neutrophil ratio across AZD1775 treatment doses. Data are shown as mean ± SEM where applicable. Statistical significance is indicated as shown.

**Supplementary Figure 5.**
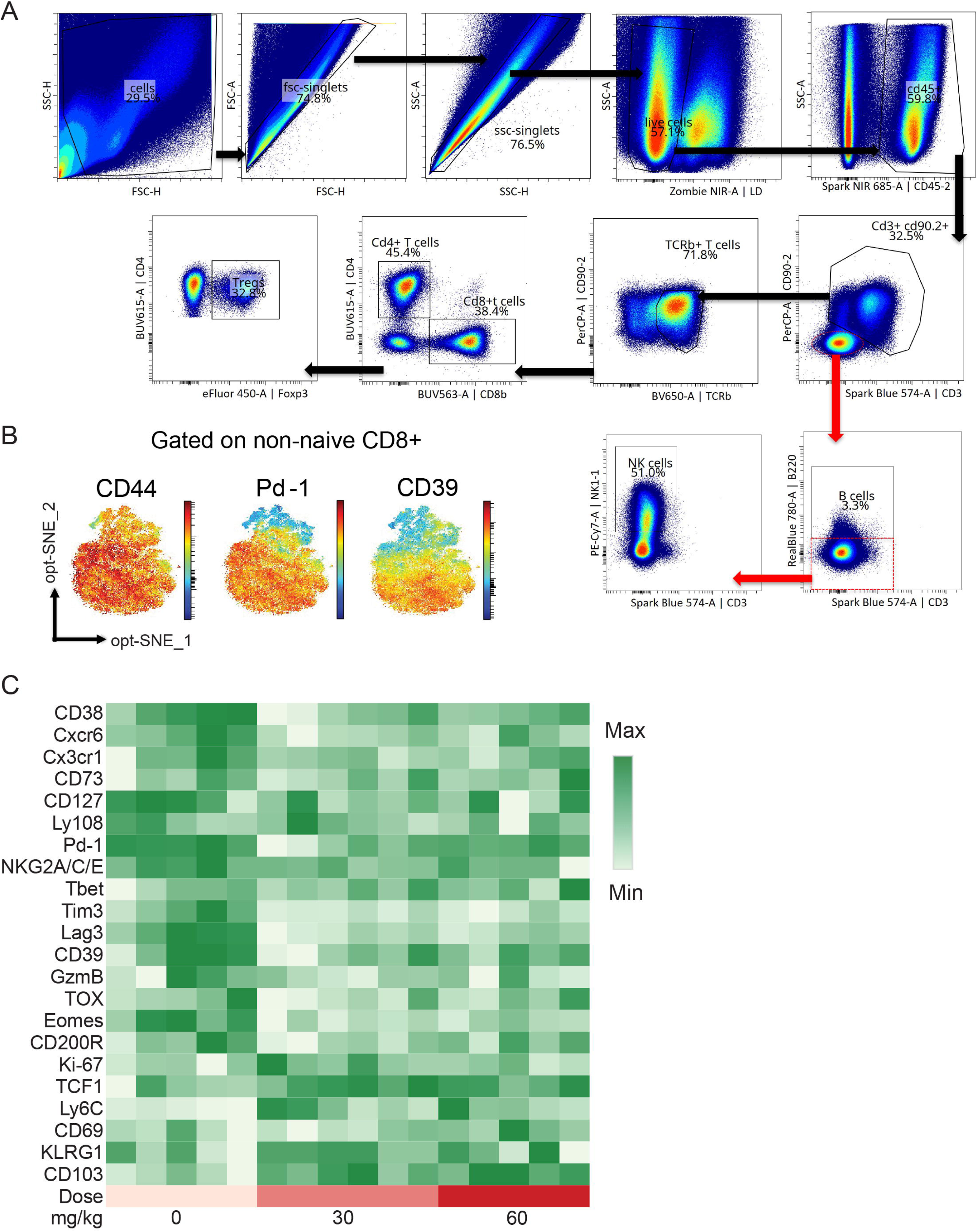
Flow cytometric gating strategy for lymphocytes in figure 5 and phenotypic profiling of intratumoral CD8⁺ T cells. (A) Concatenated scatter plots of all tumor samples (n=16) from mice in figure 5 showing gating strategy for major lymphocyte subsets. Sequential gates include exclusion of debris and doublets, selection of live cells, identification of CD45⁺ leukocytes, CD3⁺ T cells, and subsequent delineation of CD4⁺ T cells (including Foxp3⁺ regulatory T cells) and CD8⁺ T cells. Additional gates show identification of NK1.1⁺ NK cells and CD19⁺ B cells from the CD45⁺ compartment. (B) opt-SNE (optimized t-SNE) projection of non-naïve (CD44⁺) intratumoral CD8⁺ T cells, with cells colored by expression intensity of CD44, PD-1, or CD39, as indicated. (C) Heatmap showing relative expression levels of the indicated phenotypic markers across intratumoral CD8⁺ T cell populations following treatment with ZN-c3 at the indicated doses. Median fluorescence intensity (Log_10_) values are scaled per marker, with color intensity representing minimum to maximum expression across samples.

**Supplementary Figure 6.**
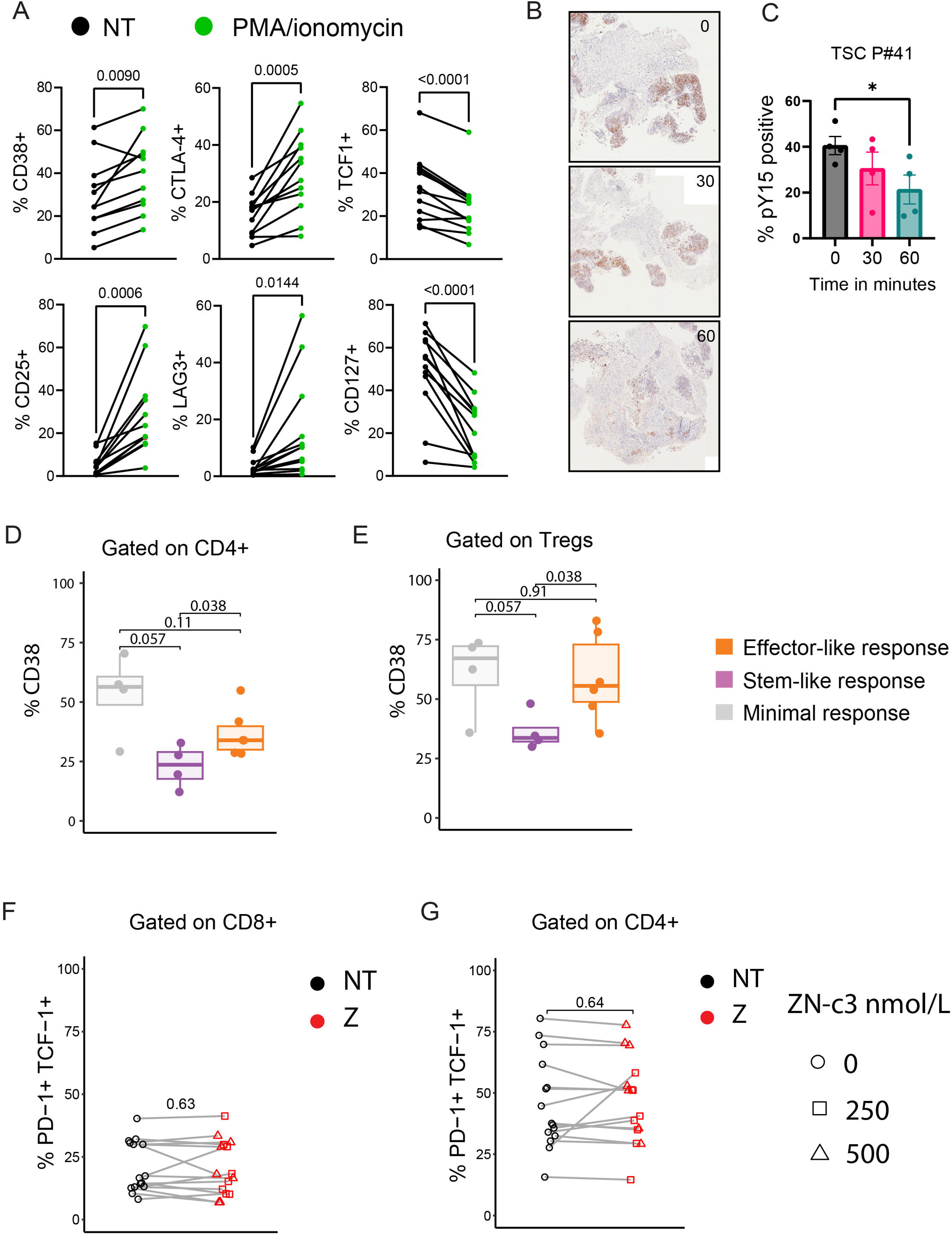
Functional phenotyping of T cells from tumor slice cultures following *ex vivo* chemical stimulation and short-term ZN-c3 exposure. (A) Flow cytometric analysis of activation and differentiation markers on T cells from human tumor slice cultures (TSC) following *ex vivo* stimulation with PMA/ionomycin for 48 hours. Paired analysis shows changes in the frequency of CD38⁺, CTLA-4⁺, CD25⁺, LAG-3⁺, TCF1⁺, and CD127⁺ subsets, relative to unstimulated (NT) controls. Each line represents a matched sample. (B) Representative immunohistochemical staining for pCDK1^Y15^ in TSCs following treatment with ZN-c3 for the indicated durations. (C) Quantification of pCDK1^Y15+^ tumor area in TSC P#41 across treatment time points. Data are shown as mean ± SEM. Statistical significance is indicated as shown. (D) Frequency of CD38⁺ cells among CD4⁺ T cells stratified by response group (effector-like, stem-like, or minimal), as defined elsewhere. (E) Frequency of CD38⁺ cells among regulatory T cells (Tregs) stratified by response group. (F-G) Paired analysis of PD-1⁺TCF1⁺ cells among CD8⁺ T cells (F) or CD4⁺ T cells (G) from human tumor slices following *ex vivo* treatment with ZN-c3 at the indicated concentrations compared with untreated controls (NT). Data are shown as individual data points with summary statistics as indicated. Statistical significance was assessed using paired or unpaired tests, as appropriate; exact P values are shown where indicated.

**Supplementary Figure 7.**
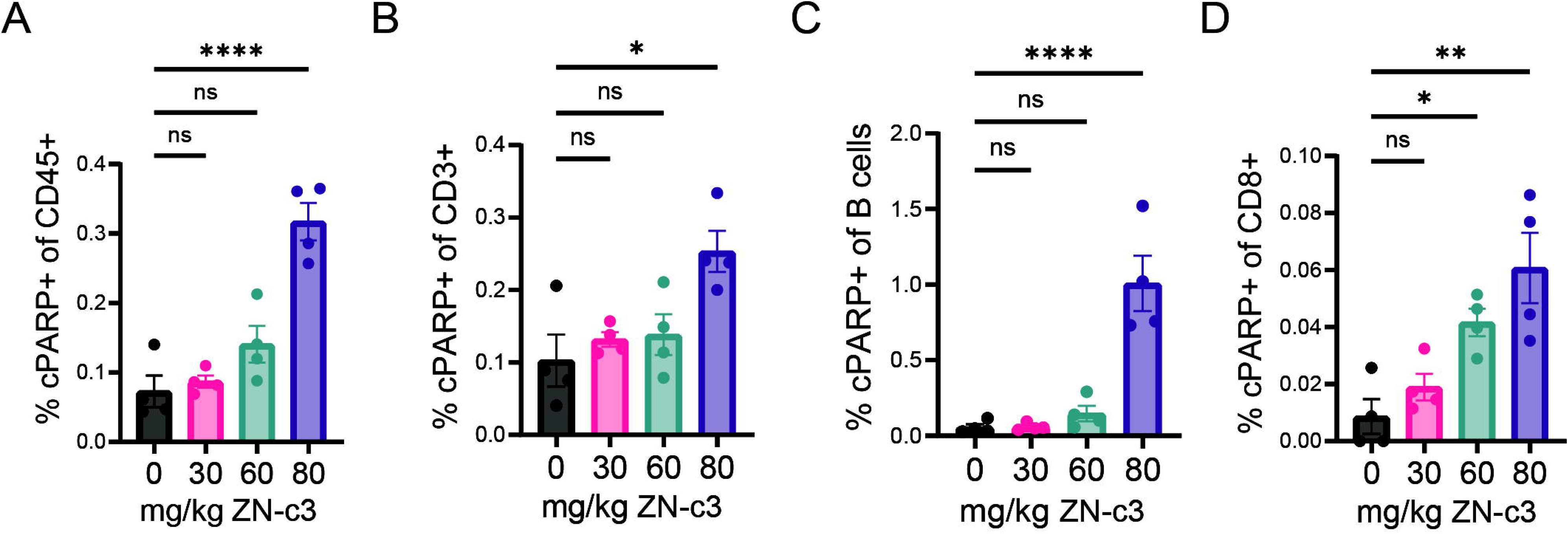
Cleaved PARP measurements in circulating immune cell populations following ZN-c3 treatment. (A) Frequency of cleaved PARP (cPARP)–positive cells among circulating CD45⁺ leukocytes following ZN-c3 treatment. (B) Frequency of cPARP⁺ cells within circulating CD3⁺ T cells across increasing ZN-c3 doses. (C) Frequency of cPARP⁺ cells within circulating B cells following ZN-c3 treatment. (D) Frequency of cPARP⁺ cells within circulating CD8⁺ T cells across ZN-c3 dosing schedules. Data are shown as mean ± SEM, with individual data points representing biological replicates. Statistical significance was determined as indicated; \**, P < 0.05; **, P < 0.01; ****, P < 0.0001;* ns, not significant.

**Supplementary Figure 8.**
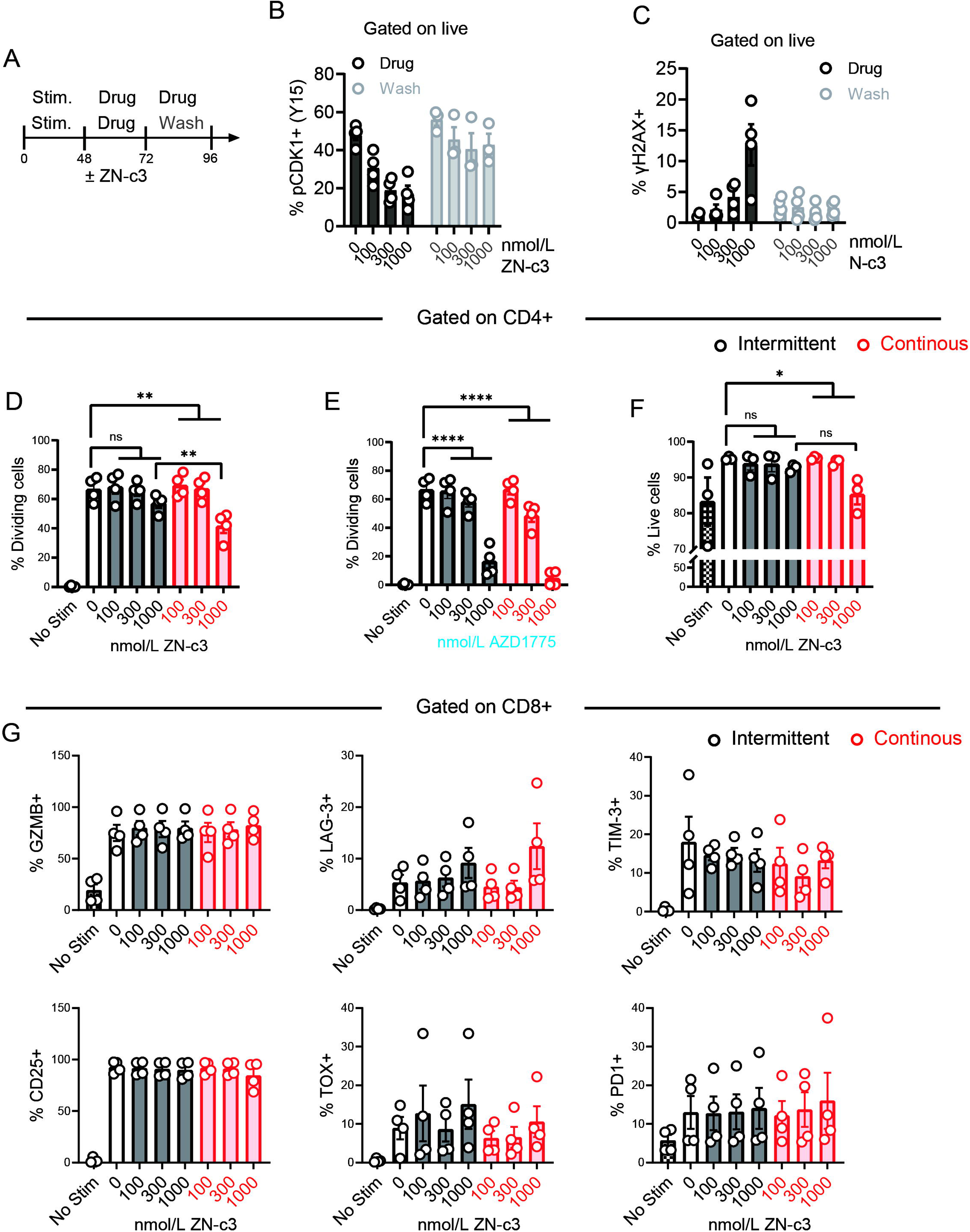
Extended analysis of the effects of intermittent and continuous WEE1 inhibitor exposure on T cell signaling, proliferation, and phenotype *in vitro*. (A) Experimental schematic illustrating *in vitro* stimulation of T cells followed by intermittent ZN-c3 exposure with washout or continuous exposure, as indicated. (B) Frequency of pCDK1^Y15^⁺ cells among live cells following ZN-c3 treatment and washout at the indicated concentrations. (C) Frequency of γH2AX⁺ cells among live cells following ZN-c3 treatment and washout at the indicated concentrations. (D) Frequency of dividing CD4⁺ T cells following intermittent or continuous exposure to ZN-c3, assessed by CellTrace Violet dilution. (E) Frequency of dividing CD4⁺ T cells following intermittent or continuous exposure to AZD1775. (F) Frequency of live CD4⁺ T cells following intermittent or continuous exposure to ZN-c3. (G) Frequencies of granzyme B–positive (GzmB⁺), Lag3⁺, Tim3⁺, CD25⁺, TOX⁺, and PD-1⁺ cells among CD8⁺ T cells following intermittent or continuous exposure to ZN-c3. Data are shown as mean ± SEM with individual data points representing biological replicates. Statistical significance was assessed using appropriate multiple-comparison tests as indicated; *P < 0.05, **P < 0.01, ***P < 0.001, ****P < 0.0001; ns, not significant.

**Supplementary Figure 9.**
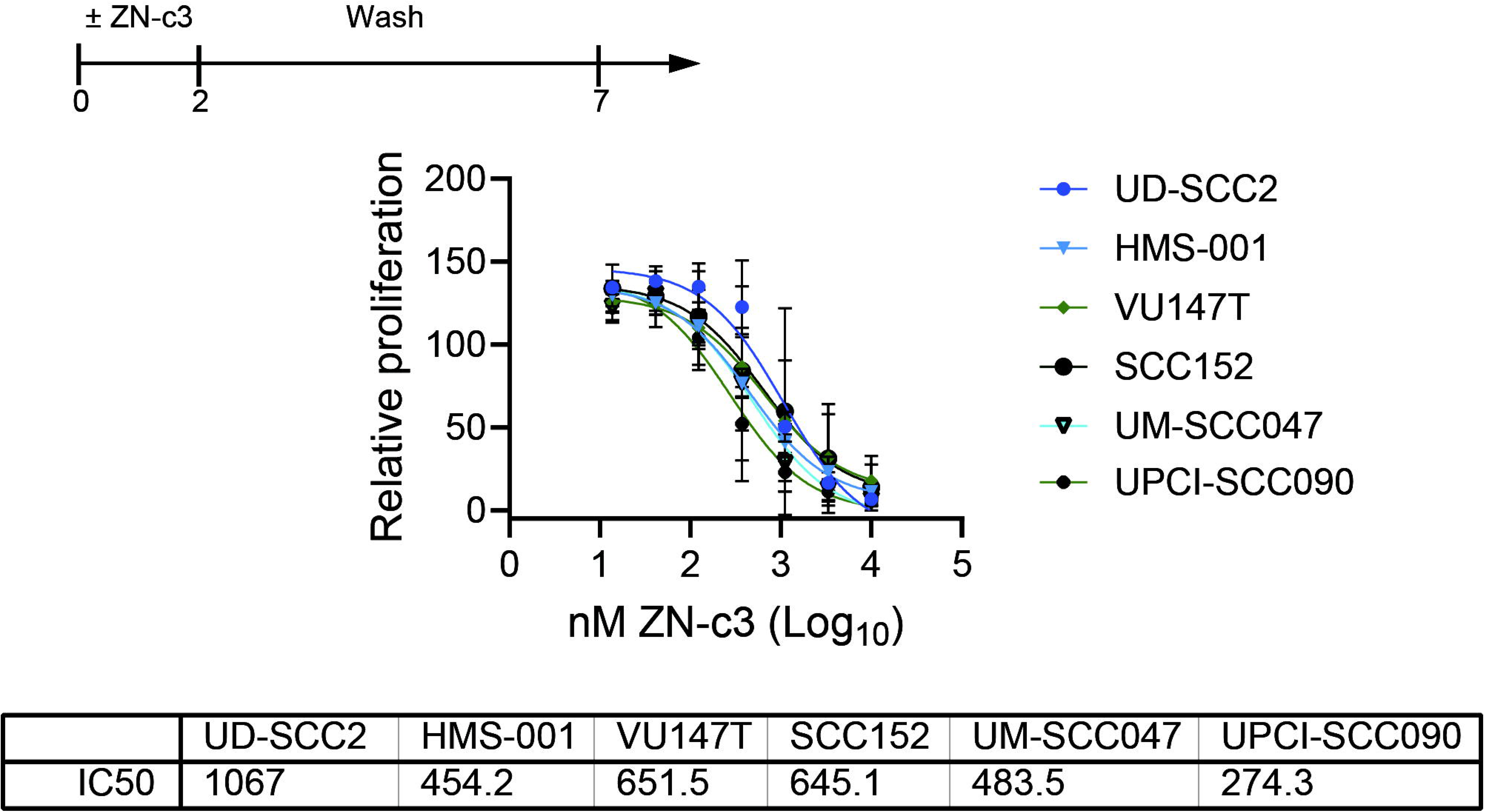
Proliferation of HPV⁺ head and neck cancer cell lines following short-term ZN-c3 exposure and drug washout. Experimental schematic depicting short-term exposure of HPV⁺ human head and neck cell lines to ZN-c3 followed by drug washout and recovery. Cells were treated with ZN-c3 for 48 hours, washed, and cultured in drug-free media until day 7. Dose–response curves showing relative proliferation following ZN-c3 treatment and washout, plotted as a function of ZN-c3 concentration (log₁₀ scale). Data are normalized to vehicle-treated controls. Table summarizing IC₅₀ values for ZN-c3 in each cell line under washout conditions. Data are shown as mean ± SEM.

