## Supplementary Methods for "Therapeutic scheduling of WEE1 inhibition preserves T cell function and promotes immune control of HPV⁺ tumors"

### **IHC Staining and Multiplex Immunofluorescence Analysis**

Multiplex immunohistochemistry and multispectral imaging were performed on 4-μm FFPE tumor sections using Opal fluorescent IHC reagents and Vectra-3 multispectral imaging systems. Slides were processed on a Leica BOND Rx stainer with sequential antigen retrieval, antibody stripping, and tyramide signal amplification to enable up to six markers per section. Primary antibodies targeted cell-cycle, DNA damage, tumor, and immune markers (Supplementary Table 7, PMID: 31852845). Images were acquired from ≥20 unbiased regions per slide and spectrally unmixed using InForm software. Cell segmentation, phenotyping, and spatial analyses were performed in HALO, with cell densities expressed as cells/mm² tissue.

### **Sample preparation for flow cytometry**

For peripheral blood analysis, blood was collected by submandibular bleed or by cardiac puncture at experimental endpoint. Red blood cells were lysed using ACK lysis buffer, followed by centrifugation and washing with PBS. If residual red blood cells were observed, a second lysis step was performed. The final cell pellet was resuspended in PBS for downstream staining.

For tumor-infiltrating leukocyte isolation, tumors were divided, with a portion fixed for histological analysis and the remaining tissue processed for flow cytometry. Tumor tissue was minced and enzymatically digested in RPMI media containing Liberase TM (1 mg/mL) and DNase I (0.1 mg/mL) for 20 minutes at 37°C with agitation. Following digestion, samples were filtered through 70-μm cell strainers, centrifuged, and resuspended in R10 (RPMI 1640 supplemented with 10% FBS; R10) media. Cell suspensions were kept on ice until further processing.

Tumor slice cultures were pooled and dissociated enzymatically using the same digestion protocol as described for tumor-infiltrating leukocytes.

### **qPCR Analysis for Gene Expression**

Total RNA was extracted from cultured cells or tissues using RNeasy Kit (Qiagen). cDNA was synthesized using SuperScript IV Reverse Transcriptase (Life Technologies) or iScript Reverse Transcriptase (Bio-Rad) following the manufacturer's protocol. Quantitative PCR was performed using PowerUp SYBR Green Master Mix (Applied Biosystems) in a 10 μL reaction volume containing 10 ng cDNA and gene-specific primers. Reactions were run on a real-time PCR system with the following cycling conditions: 50°C for 2 minutes, 95°C for 2 minutes, followed by 40 cycles of 95°C for 15 seconds and 58°C for 15 seconds. Primers are listed in Supplementary Table 8. Melt curve analysis was performed to verify amplification specificity. Relative expression was calculated using the ΔΔCt method, and results were normalized to expression of the reference gene (encoding β-Actin).

### **Immunoblotting**

Dissociated cells were washed in PBS and resuspended directly in 1X Bolt LDS Sample Buffer and denatured for 30 minutes at 95°C to obtain whole cell lysates. Lysates were separated on 4%-12% gradient gels at 80 to 150 V, and transferred to nitrocellulose or PVDF membranes using the Trans-Blot Turbo system (Bio-Rad). Membranes were blocked in 5% non-fat milk in TBST at room temperature for 1 hour, then probed with primary antibodies overnight at 4°C. The following day, after 3 washes in TBST, membranes were incubated with HRP-conjugated secondary antibodies (1 to 2 hours, room temperature). Antibodies are listed in Supplementary Table 9. Proteins were visualized using Clarity ECL substrate (Bio-Rad #1705062) and imaged on an Azure Biosystems c600 machine.

### **Cytokine Arrays**

Culture supernatants from mEERL and mTE cells treated with ZN-c3 (500 nmol/L, 72 hours) were collected. Conditioned media were spun at 1,500 rpm to remove debris, and the supernatant was stored at −80°C. Conditioned media were analyzed using the Mouse Cytokine Array AAM-CYT-3 (R&D Systems) according to the manufacturer's instructions. Quantitation was performed using the protein array analyzer macro in FIJI. Measurements were performed in duplicate.

### **NanoString Assay**

Total RNA was extracted from frozen tumors collected 24 hours after the last dose of treatment using the RNeasy Midi Kit (Qiagen). or according to the manufacturer's instructions. After quantification of RNA concentration and purity using a Thermo Fisher Scientific NanoDrop, 100 ng of total RNA for each sample was analyzed using the nCounter technology and the nCounter Mouse PanCancer Immune Profiling Panel (cat # XT_PGX_MmV1_CancerImm_CSO) according to the manufacturer's instructions (NanoString). Data normalization, differential expression analysis, and pathway enrichment (using gene set variation analysis, GSVA) were performed using NanoString nSolver Analysis Software v3.0 or Rosalind Platform (Rosalind, Inc.).
